# Stone Age anthropogenic impacts to forest development in the interior Scandinavian Peninsula

**DOI:** 10.1101/2025.01.15.632029

**Authors:** Anastasia Bertheussen, David K. Wright, Svein Olaf Dahl, Jago J. Birk, Jonas Bergman, Joseph Buckby, Sabine Fiedler, Axel Mjærum

## Abstract

Forest growth and development is a highly studied phenomenon in which humans have proven to be influential in shaping structure and composition. Delineating the long-term processes of human-environment interactions is crucial for understanding the history and trajectory of landscape formation and vegetation development. Yet, extensive knowledge of the ecological impacts of Stone Age anthropogenic activity is still lacking, particularly from Fennoscandian sites. A sediment core from South Mesna lake from the interior Scandinavian Peninsula was extracted to investigate the long-term evolutionary effects of human-environment interactions following deglaciation (c. 10,500 cal. BP) and initial colonization of the region. Analysis of the core involved microscopic/trace analytical methods, including geochemical analysis, stable isotope analysis, fecal biomarker analysis, and pollen analysis. The combined evidence demonstrates that anthropogenic impacts are prominent shapers of the ecological trajectory and landscape development of the region since the Early Neolithic (c. 5900 cal. BP), which has left a footprint on modern-day land cover.

**Significance:** Anthropogenic impacts on the environment have been observed to have had notable consequences on global and regional ecological trajectories and environmental development. Yet, how coupled human-environment interactions affect the long-term ecological complexion of boreal landscapes, such as those of the interior Scandinavian Peninsula, is not widely studied. This article presents evidence of early anthropogenic impacts to a forest’s ecology during the Holocene. Fecal biomarker and pollen analyses make it possible to provide micro-archaeological evidence of human activity and correlate it with noteworthy changes in forest structure. Our data points to a specific co-evolutionary forest development trajectory which is connected to millennial-scale human settlement patterns.

---

It is increasingly recognized that modern environments are shaped by past human-nature interactions, and anthropogenic variables must be considered to understand changes in landscape development and ecological patterns (cf. 1, 2). Advancements in paleolimnology enhance our ability to observe anthropogenic impacts in individual catchments over long timespans (cf. 3, 4). The development of novel methodological approaches in archaeology and paleoecology has provided further opportunities to investigate these effects and augmented our understanding of human-environment dynamics (cf. 5, 6, 7). However, despite increased research efforts, the long-term effects of prehistoric anthropogenic influence on the landscape development of boreal regions, such as the interior Scandinavian Peninsula, are not extensively known.

Using novel methods such as fecal biomarker analysis combined with traditional paleolimnological analyses, we can obtain evidence of prehistoric human activity in an area with poor archaeological resolution (8, 9). The use of fecal biomarkers has proven effective at reconstructing animal and human ecologies at deep time scales (9, 10). Fecal biomarkers are molecular fossils derived from different sterols metabolized through animals’ digestive systems (11, 12). Different diets and metabolic processes result in distinctive compound patterns among different species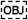(13). Humans’ digestive systems and gut microbial composition excrete high quantities of the stanol known as *coprostanol* (5β-cholestan-3β-ol) relative to other animals (8, 14, 15). Coprostanol is, hence, a prevalent marker in human feces and can consequently be used to investigate relative population dynamics over time at local scales (cf. 9, 14, 16). In this paper, we use coprostanol in parallel with other environmental data such as stable isotopes (δ^13^C and δ^15^N) from terrigenous organic matter and pollen to examine the long-term effects of human activity on the growth and development of a boreal forest in the interior Scandinavian Peninsula.

## Study Site

South Mesna lake (SML) is found in the county of Innlandet, Norway (Fig. 1) at approx. 520 m.a.s.l., and is part of the Mesna drainage system which connects to the larger drainage system of Gudbrandsdalslågen. The area is primarily underlain by sandstones (greywackes and arkose), shales and conglomerates (17, 18). Glacial deposits dominate the near-surface geology, including downhill eskers, kettle holes, and hummocky moraines (17). A recent study by Mangerud et al. (17) revealed that the local area was deglaciated around 10,500 cal. BP where birch (*Betula* spp.) formed the first woodland. The introduction of Scots pine (*Pinus sylvestris)* and the creation of boreal forest likely did not occur until after 10,000 cal. BP (also 19, 20). However, during the first two thousand years after deglaciation forest development fluctuated, and varying amounts of sedges (*Cyperaceae)* and grasses (Poaceae), indicating the opening of the local forest, could be identified in the region (17, 20).

**Figure 1.**
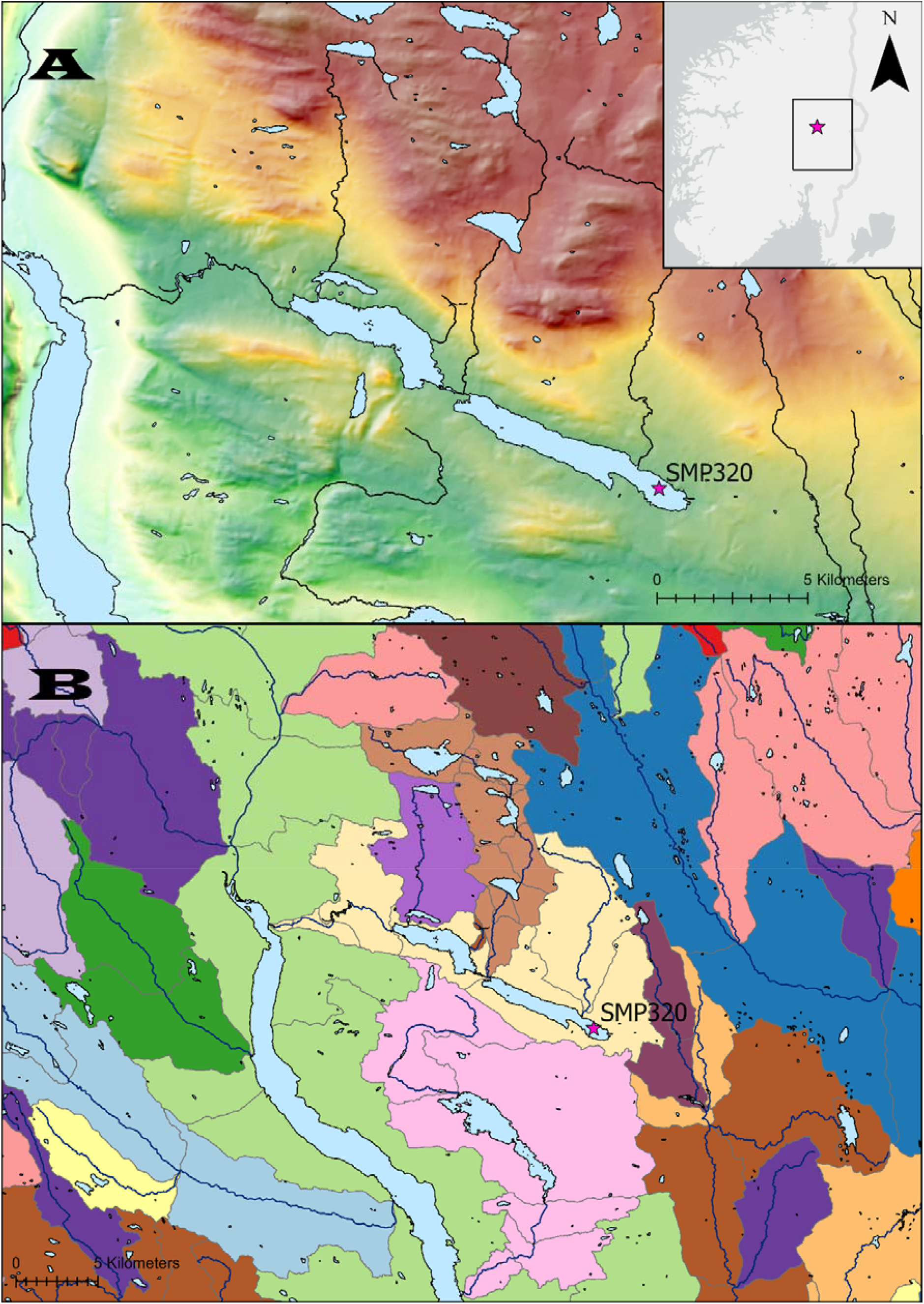
A) Topographic map of the study area B) A watershed map displaying the catchment areas. Data gathered from The Norwegian Water Resources and Energy Directorate (NVE) and the Norwegian Mapping Authority (Kartverket).

Shortly after deglaciation, hunter-gatherers in Fennoscandia are known to have traversed both coastal and inland landscapes (cf. 21). Yet, due to survey bias, there are fewer archaeological data from the interior Scandinavian Peninsula compared to the coast. However, subsistence strategies among coastal and inland hunter-gatherers appear to be comparable, with seasonal fishing, hunting, and plant collection occurring in both groups though there are significant variations in toolkits, use of raw materials and targeted prey (22, 23). Nonetheless, interior settlements, especially the ones situated adjacent to waterways and geographically closer to the coast shared many cultural traditions and are deemed similar enough to be comparable to coastal sites in socio-cultural development (18, 22). Even so, we have an incomplete understanding of Mesolithic hunter-gatherer lifeways and their landscape utilization practices in the interior region.

A total of 144 archaeological objects have been heretofore identified in the vicinity of SML and the neighboring North Mesna Lake through either stray find donations or archaeological registrations (24)(Fig. 2). These are too few archaeological objects to paint a clear portrait of the region’s settlement history. Yet, these findings strongly support the hypothesis of a human Mesolithic presence in the area. Some objects, such as microblades and microblade cores, could potentially have been produced during the period immediately following deglaciation (10,250–8250 cal. BP), while the presence of oblique arrowheads indicates human activity around the Mesolithic/Neolithic transition (c. 5900 cal. BP) (25). Human presence in the early and middle Neolithic period (c. 5900-4400 cal. BP) is clearly evidenced by the discovery of slate arrowheads, part of a polished flint axe of South Scandinavian type, and a hearth containing elk bones directly dated to c. 5500 cal. BP (24, 26).

**Figure 2.**
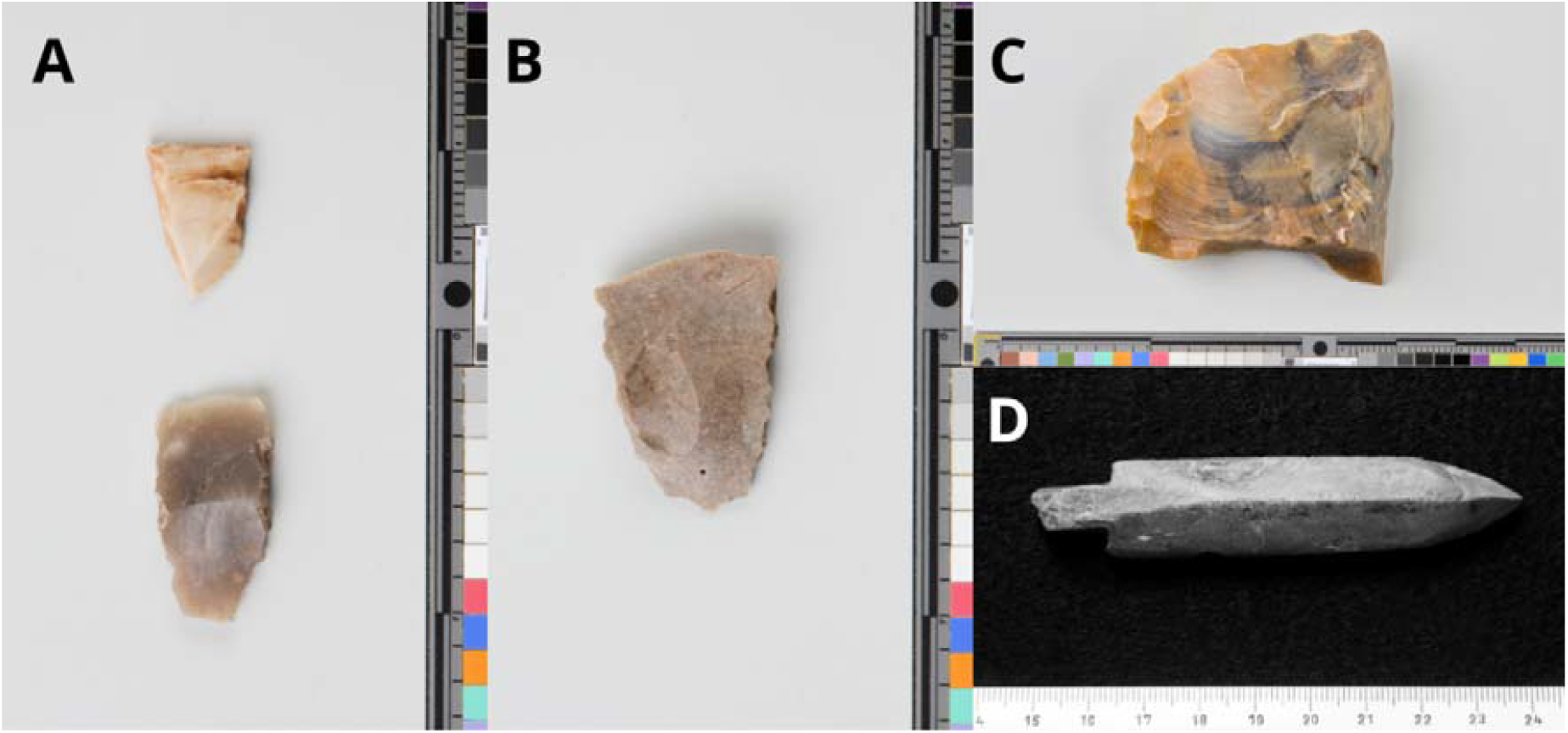
A selection of archaeological finds found along the Mesna lakes A) Flint oblique arrowheads B) Flint oblique arrowhead C) Part of a flint axe D) Slate arrowhead, photo by Ove Holst. All photos ©Museum of Cultural History, University of Oslo.

During the Neolithic in southern Norway, settlements were not necessarily more sedentary than their predecessors. It has been argued that the introduction of agriculture into the region was a long process and likely occurred multiple times (e.g., 27). Subsistence agriculture was likely not integrated before the late Neolithic (28, 29). Therefore, during much of the Neolithic, it has been presumed that landscape utilization practices had ecological impacts similar to the Mesolithic (4). Artifacts and archaeological sites dated to later periods such as the Bronze Age and Iron Age (c. 3650–900 cal. BP) indicate pulses in human activity in the area rather than the presence of sustained, long-term occupations (e.g., 18, 24, 30). Nevertheless, it has been hypothesized that low-scale agricultural or “gardening” occurred during the earlier parts of the Neolithic (29), which later gave way to more intensive and impactful forms of land appropriation. If this was the case, the introduction of land management practices such as clearing forests for hunting or small-scale plant and/or livestock husbandry may have taken place as early as initial parts of the Neolithic, altering the ecological complexion of the landscape far earlier than previously considered. To what extent such practices took place remains an open question because such effects have yet to be studied on catchment scales in paleoecological records. Our research tests this scenario with a fecal biomarker, pollen and stable isotope (δ^13^C,δ^15^N) reconstruction of a laminated lake core to understand the long-term ecological formation processes from SML.

## Results

A continuous sediment core (SMP320) was retrieved at 61.072°N, 10.819°E from the lacustrine sediments of SML during the summer of 2020 with a piston corer attached to a gravity auger fixed to a pontoon raft. From this core, we reconstruct the paleoenvironment with the main objective of identifying the impacts of human activity and anthropogenic landscape utilization on the evolution of the ecology of the catchment.

### Stratigraphic sequence

The core is comprised mainly of intercalated silty-sand and clay-rich layers (Fig. 3). The silty-sand layers consist of gyttja and organic-rich slope wash sediments. In contrast, the clay-rich layers are minerogenic and deposited as slow sedimentation during high-water episodes. Loss-on-ignition (LOI) values show significant variability in total carbon content observed where an overall higher organic matter content is detected episodically intercalated with cm-scale minerogenic laminates. The values range between ∼1 to 24% total organic carbon (TOC). Thirteen lenses in SMP320 were selected for Accelerator Mass Spectrometry (AMS) radiocarbon (^14^C) dating. Due to the prevalence of mainly aquatic plant macrofossils, only nine samples had usable terrestrial plant macrofossils for dating. The interval of the nine ^14^C samples extends between 9090 and 770 cal. BP (Supplementary table X.1). The LOI data indicates three distinct minerogenic depositional phases: 6610–6450 cal. BP, 5200 – 5000 cal. BP, and 4350 – 4300 cal. BP, with the middle episode showing the lowest values where multiple LOI samples fall below 5% TOC. An increased sedimentation rate is observed from c. 5560 until 4700 cal. BP. A mean sedimentation rate of ∼0.85 mm/yr is noted during this interval, double the average core sedimentation rate of the core of ∼0.46 mm/yr. The highest rate is observed between 5560 and 5400 cal. BP with ∼1.24mm/yr.

**Figure 3.**
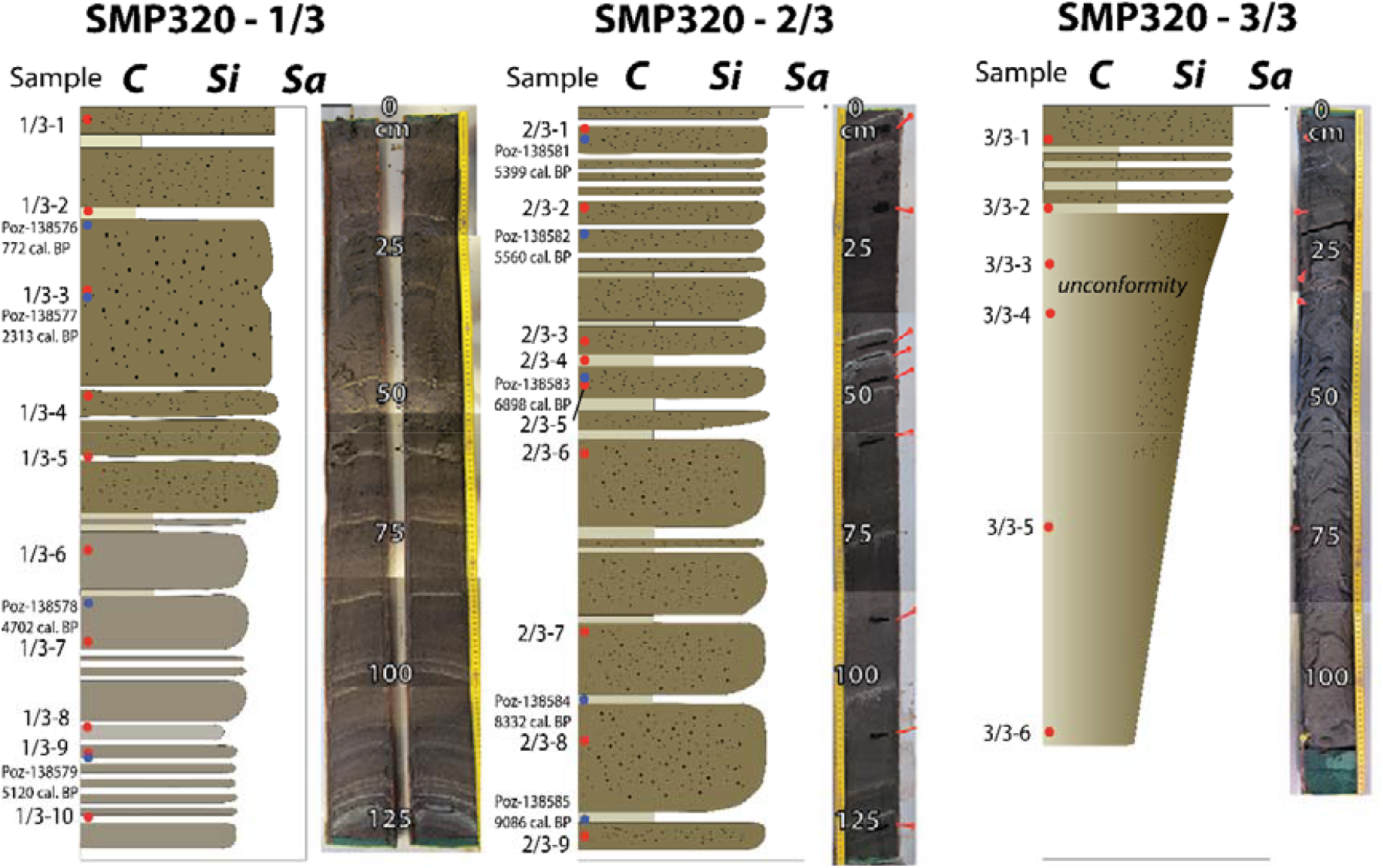
Sediment core SMP320.

### Vegetation dynamics

A primarily boreal forest environment is presently found around SML. With the stable isotope- and pollen analyses the intention is to evaluate forest development and long time-scale vegetation succession. A total of 25 sediment samples were analyzed for bulk δ^13^C and δ^15^N, while 15 samples were chosen for pollen analysis (Fig 4). The δ^13^C values range between -30.51‰ and -25.68‰ (vs. VPDB), while the δ^15^N ranges between 2.68‰ and -0.86‰ (vs. AIR). Variability in the isotopic data is observed, but primarily, a pattern of decreasing δ^13^C and increasing δ^15^N is detected. Two samples show significant deviations from this pattern. Between sample 2/3-5 and 2/3-4 the δ^13^C enrich from - 29.05‰ to -27.30‰, and between sample 2/3-1 and 1/3-10 the sample values enrich from -29.19‰ to -27.79‰. Correspondingly the δ^15^N depletes from 2.65‰ to 0.43‰ and from 2.68‰ to 0.87‰, respectively. Sample 2/3-4 is interpolated to date from the late Mesolithic (6940 – 6470 cal. BP) and sample 1/3-10 from the transition between the early Neolithic/middle Neolithic (5450 – 5090 cal. BP).

**Figure 4.**
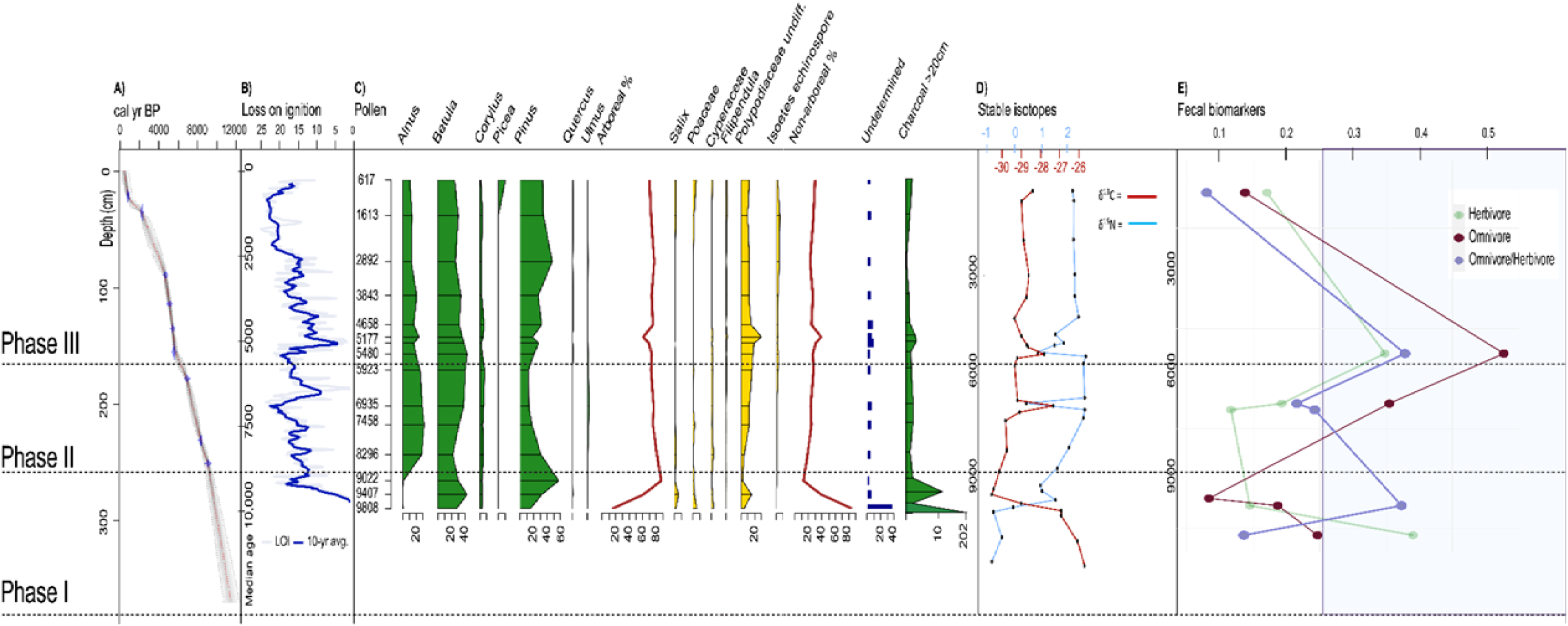
Proxy reconstruction of sediment core SMP-320 divided into three ecological phases. A) An age-depth model; B) LOI results where the dark blue is a 10-yr interval; C)Pollen analysis results of species that had a total abundance of 10% or more compared to other taxa as well as a charcoal count; D)Stable isotope results in which red is δ^13^C and the blue is δ^15^N; E) Results of the fecal biomarker analysis in which green is 5β-sitgmastanol/5α-stimgastanol (herbivore), red is (coprostanol+epi-coprostanol)/cholestanol (omnivore) and blue is coprostanol/5β-stigmastanol (omnivore/herbivore).

SML serves as a regional pollen sink because of its large size and extensive river connections. Due to their pollination dispersal patterns, trees and other wind-pollinated plants are regional vegetation markers. In contrast, herbs and insect-pollinated plants are signals dominated by the immediate area within the drainage catchment. The pollen concentration is noticeably low in the oldest sample, however the concentration throughout the remainder of the core is sufficient for vegetation reconstruction. The total sums of pollen studied in each of the 15 samples are generally between 700 and 800. A total of 59 unique taxa were identified of which 48 are terrestrial pollen types. Around 4– 10% of the pollen taxa are unidentifiable, likely reflecting the significant erosion of SML’s beach zones during deposition.

The pollen analysis reveals that between 9810–9410 cal. BP, heliophylic plants such as birch (*Betula* sp.), hazel *(Corylus* sp.), Poaceae and other grasses, sedges and sorrels dominated the landscape. Around 9410 cal. BP pine reaches its maximum percentage value of 57%, reflecting its regional dominance. Alder increases sharply from 9020 cal. BP where it only represents ∼1% of the assemblage to ∼28% around 8300 cal. BP. This expansion continues until around 5480 cal. BP, where alder becomes a prevailing genus. Towards the end of this period hazel reaches its highest presence in tandem with other deciduous trees such as elm (*Ulmus sp*.), linden (*Tilia* sp.), and willow (*Salix* sp.), which also increase. Charcoal does not vary significantly during this period. From 5940 – 1680 cal. BP, hazel decreases from representing ∼6% of the pollen sum to only ∼3%. Yet, other deciduous trees such as oak (*Quercus* sp.) and lindens increase during this period, and ash trees (*Fraxinus excelsior*) are observed for the first time around 5460 cal. BP. Between 5360 – 5000 cal. BP charcoal quantities more than double from 1.97% (charcoal particles ≥20 µm) to 4.22%. This corresponds to an increase in ferns (family Polypodiaceae), from representing 17% of the pollen assemblage to 30% around 5000 cal. BP. Shrubs and herbs slightly increase around 3390 cal. BP, while closed-canopy taxa such as lindens decrease. Deciduous trees start to decrease from this period representing 69% of the tree pollen count down to 48% around 2890 cal. BP. The semi-woodland taxon *Juniperus sp*. reaches its maximum value around 1610 cal. BP, which occurs frequently as a disturbance taxon in pastures (6). Clear traces of pastures and cultivation are indicated around 600 cal. BP on the basis of taxa such as barley (*Hordeum* sp.) as well as high grass counts.

### Fecal biomarkers

The highest concentrations of steroids recovered in the SML core derive from plant membranes classified as phytosterols, such as β-sitosterol and stigmasterol, as well as cholesterol from animal tissue. Concentrations of β-sitosterol is ≤322 µg/g_TOC_, stigmasterol is ≤59 µg/g_TOC_ and cholesterol is ≤69 µg/g_TOC_. The highest concentration of each sterol occurs in different layers of the core. Similarly, the environmental reduction products from these compounds have also higher concentrations than fecal biomarkers. Correspondingly, the concentration of cholestanol is ≤42 µg/g_TOC_ and 5α-stigmastanol is ≤140 µg/g_TOC_. These reduction products have their highest concentrations in different samples and a common maximum is not identified at a specific depth. The fecal steroids, i.e. coprostanol and 5β-stigmastanol (as well as their 3α-epimers epi-coprostanol and epi-5β-stigmastanol) are identified in varying concentrations throughout the core, but in significantly lower quantities than the Δ^5^-steroids and 5α-stanols. Coprostanol has concentrations of ≤13 µg/g_TOC_, 5β-stigmastanol have concentrations of ≤34 µg/g_TOC_, whilst their 3α-epimers where not found in all samples and never exceed 10 µg/g_TOC_. The highest concentrations of all fecal biomarkers are observed between 5450 – 5090 cal. BP (sample 1/3-10).

We have determined ratios to correct for environmental background levels of stanols (cf. 14, 31) (3α-epimers where not included in these ratios because of their absence in some samples but including them into the ratios did not change the interpretation and these ratios are shown in the SI). Samples with concentrations equal to or above the median are deemed to have enhanced fecal input which includes sample 1/3-10 (c. 5450 – 5090 cal. BP), 2/3-4 (c. 6940 – 6470) and 3/3-5 (c. 11,470 – 10,060). These samples are additionally calculated to determine whether the enhanced fecal input is omnivorous or herbivorous where the median ratio is determined to be 0.23. Sample 1/3-10 has an above median ratio of 0.38, whilst 2/3-4 has a ratio of 0.22, i.e. just below the median. The oldest sample, 3/3-5, has a low ratio of 0.14. Bile acid values do not exceed environmental background levels and are therefore not incorporated into the analysis. However, the bile acid HDCA, which is most pronounced in pigs (*Sus* sp.), is not detected in any of the samples, even in trace background levels.

## Discussion

The combined geochemical and plant fossil evidence at SML indicates distinct phases of forest succession in concert with human occupation of the region. The synthesis of these results indicates that detectable canopy cover changes and an uptick in sedimentation rates correlate with evidence for human presence on the landscape vis-à-vis elevated fecal input into the system.

### Phase 1 (12,320 – 8940 cal. BP): Deglaciation and initial vegetation succession

Samples from this core section reflect the initial vegetation colonization and succession following deglaciation. They serve as an anchor reference point for the environmental processes of forest growth with limited evidence of human impact. We base this assertion on the lack of archaeological objects, from our site as well as adjacent archaeological sites in the region (e.g., 18, 32), and the coprostanol values which suggest that, if present, humans had very limited impacts on the landscape.

There is evidence for an open, yet afforesting environment following deglaciation from around 9330 cal. BP. Stable isotope analyses indicate negative δ^15^N values and isotopically enriched δ^13^C values (≥-27‰) in the deepest four samples dated to before 9330 cal. BP. The samples that overlie this basal aspect of the core reflect a steady increase in δ^15^N and a steady depletion of δ^13^C values, which we interpret as a progressively more closed-canopy forest (33, 34). Biomarkers from sample 3/3-5 indicate a higher fecal input than in the other samples (high ratios) primarily from herbivore species. This is likely due to an increase in forest-dwelling animals using the lake as a water source as biological production increase (35-37).

### Phase 2 (8940 – 5810 cal. BP): The Mesolithic entrance

A warmer and drier climate in the mid-Holocene likely facilitated more boreal forest growth (38), though a colder interval has been noted in a previous study (39). Our data also indicate continuous canopy growth into Phase 2, and increasingly depleted δ^13^C values can be detected until around 8420 cal. BP. Regional paleoclimatic studies show precipitation increased during Phase 2 (40, 41), which may have led to episodic lake overfilling. Our LOI data also indicate increased minerogenic content between 6610–6450 cal. BP which strengthens this premise.

A change in our data is detected between 6940 – 6470 cal. BP, where a significant enrichment in the isotopic values is observed, revealing a sudden change in the forest structure to a more open canopy. The results point to a “canopy effect” in which a lower contribution of ^13^C-depleted CO_2_ respired from the forest floor because more sunlight can be dispersed to shrubs and grasses, thus providing more enriched δ^13^C values (33). Concurrently, the fecal biomarkers indicate that there are more omnivorous animals present on the landscape. This, due to the lack of the HDCA bile acid and other native omnivorous species, is interpreted to be the result of more human presence around the SML catchment. Human activity is additionally corroborated through the archaeological record. Less herbivore presence could also be indirect evidence of an *ecology of fear*, i.e. that novel predator (i.e., human) habitat encroachment affected the distribution and behavior of wild animals (42). No significant increase in charcoal is noted in this phase, so change in the fire ecology is unlikely to have altered the trajectory of the forest development from this time period. However, there is a possibility that charcoal simply has not accumulated within this aspect of the lakebed.

With samples following 6470 cal. BP, an increase in δ^15^N values indicate a decrease in canopy-cover with accelerated oxidization of NH_4_^+^ into NO _3_^-^ in open environments, which increases the plant-available N pool and therefore δ^15^N enriches within the understory plants and surface soils. This is additionally supported by the increase of heliophylic plants such as heather (*Calluna vulgaris)* in the pollen assemblage. Possibly due to warmer temperatures, indicated by the δ^15^ N values, a natural “shrubification” could have been occuring. Since a climatic change has been noted for this region previously (cf. 39), we interpret the abrupt canopy cover change as a likely expression of the climatic variability from this period. However, we acknowledge the possibility of anthropogenic activity as supported by the archaeological record (4). Moreover, an increase in hazelnut vegetation is observed between 6460–5650 cal. BP. Hazelnut trees have long been associated with human activity in the Scandinavian Peninsula, as the species does not possess seeds that are easily transported by wind, which makes it reliant on animals for dispersion (43, 44). Hazelnut trees commonly thrive better in a more open canopy as it is an understory species (43, 44).

### Phase 3 (5810 cal. BP – present day): An anthropogenic ecological trajectory

An overall regional cooling trend has been previously identified between 6500 – 5000 cal. BP (39, 45). However, it has also been argued that during this part of the middle Holocene southern Norway had stable, mild temperatures comparable to present conditions (46). Nevertheless, the conditions were cooler than the previous warm conditions of 1-2°C above present temperatures, which could have led to an increase in effective moisture (40). We observe a doubling of the sedimentation rate in our age-depth model beginning between 5560 and 5400 cal. BP, indicating increased erosion within the basin. This could reflect increased effective moisture combined with fewer trees on the hillslopes to restrict solifluction. However, the δ^13^C show depleting values which can be interpreted as an initial increase in canopy-cover.

Around 5450–5090 cal. BP enriched δ^13^C values are observed indicating more open land cover conditions. The pollen data also corroborates the presence of more open canopy through increased ferns and grasses whilst all deciduous trees decrease. Simultaneously, the LOI values change from ∼20% to ∼3% which can be explained by less vegetation cover and, thus, less influx of organic matter. Concurrently, coprostanol, i.e., the human fecal biomarker, is observed to have its highest concentration measured from the core and the stanol ratios indicate enhanced input of omnivorous feces. Early to middle Neolithic (c. 5850 – 4300 cal. BP) archaeological objects such as slate arrowheads additionally support the biomolecular evidence for an uptick in human activity.

Furthermore, we observe an increase in charcoal which is interpreted to correspond to a higher frequency of fires to open and maintain clearances for subsistence pursuits. This is also corroborated with a slight uptick in heather. Heather is a disturbance plant that thrives after burning and often expands following anthropogenic clearances (47). Post-burn heater shoots are also argued to be more nutritious for grazing animals than older ones, providing an incentive for pastoralists to practice pyroculture (48). Although traditional large-scale agricultural practices were not common in the boreal landscapes during the early/middle Neolithic in Fennoscandia, the use of both forest grazing and clearance fires for small-scale agro-pastoralism has precedent in the anthropological record. The use of fire as a landscape management tool by hunter-gatherers and small-scale farmers is well-known globally from ethnographic sources (49-51). The use of fire leads to the creation of forest mosaics, which have been proven to encourage species variation and hence improve resource accessibility (51). However, studies from southern Scandinavia indicate that small long-lasting clearances for grazing cattle were more commonly created than the utilization of the slash-and-burn technique during the earlier parts of the Neolithic (52). We therefore argue that the observed increase in charcoal is a reflection of the intentional creation of open patches in the forest for agropastoral pursuits. Additionally, omnivorous presence is the highest, but herbivore values also increase during the same period, which could be due to an increase in domestic animals. We, therefore, argue that human land management practices impacted the evolution of this northern boreal landscape before the introduction of full-scale agriculture to the region.

Following this period of inferred anthropogenic canopy cover change, i.e., in samples younger than 5250 cal. BP, the stable isotope values never return to the environmental baseline observed in previous sections. A decline in thermophilic vegetation is observed from around 5200 cal. BP which might indicate a change to a colder climate which corresponds with past temperature reconstructions that argue for a cooling from around 5000 cal. BP (40, 53). From 4120–2950 cal. BP a more persistent human forest management strategy is detected as the δ^13^C values stabilize above -29‰. The prevalence of sedges, grasses, and heather also increases, while linden decreases, which signals that anthropophilic meadowlands and pastures have become permanent features of the landscape. The sole biomarker sample is from c. 430 cal. BP where the biomarker samples are lower than the previous. However, their median is still high enough to be indicative of continued activity. Although the human activity during this period was likely concentrated in other parts of the lake with sediment sinks different than the SMP320 core. For example, in the southern end of North Mesna lake, a fishing trap was excavated and dendrochronology dated to the same time period, i.e. the 14^th^ century (54). Additionally, one pollen sample dated c. 590 cal. BP from the area indicates plant cultivation was prevalent around the SML with the identification of barley (*Hordeum vulgare*). Rye (*Secale cereale L*.) was also found in the low-magnification analysis. Furthermore, the herbivore signal is higher than the omnivore signal which could be a shift to different animal husbandry strategies i.e. more pastures or a change of animals. However, this has to be further evaluated as our evidence is inconclusive.

Given the sum of the results, we argue that there is a direct and profound change to the forest’s evolutionary path following Early Neolithic expansion into the region where vegetation cover remained more open than climate conditions predicted (33, 40). Other regional vegetation studies similarly argue that farming in the late Neolithic and Bronze Age on the Scandinavian Peninsula shaped mosaic landscapes where pastures were adjacent to forests (43). We therefore argue that the contemporary landscape of SML has developed with the active intervention of humans, which has had notable signficant impacts on the ecological trajectory of the forest.

## Conclusion

In this study, we identify alterations in the functional ecology of SML beginning around 5200 cal. BP, when post-glacial forest succession gave way to more open, fire-prone conditions at the advent of the Neolithic period in southern Norway. Through the use of a multiproxy methodology, specifically the use of micro-archaeological techniques, it has been possible to observe anthropogenic impact to a paleoenvironment in a landscape that has been severely affected by Late Holocene erosion. Our study suggests that Mesolithic foragers likely did not affect the ecosystem function as drastically as natural climate variability. However, with an opening of the forest canopy cover, an increase in heliophylic plants, and an increase in coprostanol, already in the initial stages of the Neolithic does anthropogenic activity appear to have had a substantial effect on the environment. Arguably this is the start of a long chain of human impacts, which formed the contemporary environment as we know it today. Ultimately, tracing the formation of early anthromes in boreal ecosystems is key to understanding how human impact affects long time-scale forest development.

## Materials and Methods

In 2020 a sediment core (SMP 320) was retrieved from South Mesna lake with a piston corer attached to a gravity auger on a pontoon raft (Supplementary text S1). The core was divided into three subsections (1/3, 2/3, 3/3) and then marked, described, and photographed (Figure 3). Nine terrestrial plant macrofossils were retrieved for AMS ^14^C (Supplementary table S2). The radiocarbon samples were extracted from the core by members of the Department of Geography at the University of Bergen and run at Poznan Radiocarbon Laboratory in Poland. The results are calibrated using OxCal with the IntCAL20 calibration curve. The dates are statistically interpolated in an age-depth model in the statistical programming language R 4.3.1 (55) using the package *rbacon 3*.*2*.*0* (56). All samples are age determined from this model, and all date ranges are presented at the 2-σ confidence level.

### Geochemical analysis

The core was marked, split, cleaned, described and analyzed at the Department of Geography/EarthLab at the Department of Earth Sciences at the University of Bergen, then transported to the Culture History Museum at the University of Oslo for further sedimentologic analysis. Magnetic susceptibility (MS), loss-on-ignition (LOI), and dry bulk density (DBD) were conducted on the core for efficient and reliable identification of the core whilst working. LOI samples were taken per 0.5 cm and dried at 550°C for one hour. In 5 cm intervals select samples were chosen to be dried additionally at 950°C to infer the carbonate content based on the LOI. Sediment DNA was also attempted but did not provide any significant results. A full geochemical report is found in the supplementary material (Supplementary text S1).

### Fecal biomarker methods

Seven samples were analyzed for fecal biomarkers: 5α- and 5β-stanols, as well as Δ ^5^-sterols concentrations. Seven samples from SMP320 and one from the modern beach mud at the landing dock of the research raft were sent for fecal biomarker analysis to the Institute of Geography at the University of Mainz. The samples were dried at 40 °C, sieved at ≤2 mm and finely ground. The method from Birk et al. (31) was used for the extraction, purification, and measurement of Δ^5^-sterols and stanols as well as bile- and fatty acids. A full biomarker report is found in the supplementary material (Supplementary text S2).

### Stable Isotope Analysis

A total of 25 samples were removed from SMP-320 for stable isotope analysis. Samples were initially prepared by separating nitrogen from carbon assays following manual pulverization. The carbon assays were agitated in 1M HCl on a Vortex shaker for 24h, rinsed and centrifuged 3 times with distilled water, dried at 40°C for 72 hours. Following this, both sets of samples were loaded into tin capsules weighed on a microbalance to 25±1 mg, which was adjusted in subsequent runs to accommodate TOC. Prepared samples were analyzed using a DeltaV Advantage Stable Isotope Mass Spectrometer configured with a Flash Elemental Analyzer (ThermoFisher) for combustion into purified CO_2_ and N_2_ at the CLimate Interpretation of Plant Tissue lab in the Department of Biosciences at the University of Oslo. The isotopes were used to enable inferences concerning changes in forest canopy cover. Varying root- and leaf litter composition and differing photosynthetic pathways garner specific isotopic values in the soil (57-60). In boreal environments, sunshine is presumed to be the primary climatic control, as opposed to extreme moisture stress, common in other biomes (33). δ^13^C_VPDB_ internal references and quality assurance calibrated against NBS19 and LSVEC with consensus values 1.95‰ and -46.6‰, respectively. δ^15^N_AIR_ internal references and quality assurance calibrated against USGS40 and USGS41 with consensus values -4.52‰ and 47.57‰, respectively (Supplementary text S3).

### Pollen analysis

A total of 15 sediment samples of ∼1 cm^3^ in size from approximately every 20 cm between 230 cm and 10 cm were obtained from the sediment core for pollen analysis. The pollen samples were enriched at the microfossil laboratory at The Archaeologists, National Historical Museums, Stockholm, according to the standard methods described by Berglund & Ralska-Jasiewiczowa (61). Samples were dispersed in sodium hydroxide (NaOH), calcium carbonate (CaCO_3_) was removed with hydrochloric acid (HCl), and cellulose was removed with acetolysis solution consisting of concentrated sulfuric acid (H_2_SO_4_) and acetic anhydride (CH_3_CO)_2_O). The samples were embedded in glycerin before microscopy. The identification of both pollen and spores was done with the help of reference literature (62) and a reference collection of modern pollen taxa. Terrestrial plants such as trees, shrubs, dwarf shrubs and herbs were added to the total pollen count, while ferns, water- and marsh plants, mosses, and algae were not included. Charcoal particles >20 µm were also counted. Low magnification analysis was also implemented to detect low-frequency pollen types at more sample levels. A full pollen report is found in the supplementary material (Supplementary text S4).

## Data and software availability

A full R script is available at DOI: 10.5281/zenodo.11487066, which can be used to reproduce the analyses and visualization. Data used can be accessed as Excel files in supplementary materials.

## Acknowledgements

This research was funded by Riksantikvaren as part of the arrangement with sector fee for heritage management in regulated watercourses. Thanks to Pål Ringkjøb Nielsen and Kristian Vasskog, Department of Geography, University of Bergen, who assisted during the coring of the lake, the technical assistant Stefanie Klassen at the Johannes Gutenberg University Mainz for the analysis of the biomarkers, and to Claudia Tamara for preparing and William Hagopian at the CLIPT Stable Isotope Laboratory at the University of Oslo for analyzing the isotope samples.

