## Supplementary data - text, tables, figures for "Stone Age anthropogenic impacts to forest development in the interior Scandinavian Peninsula"

**This PDF file includes:**

**Supporting text (Text S1–S4)**

**Figures S1 to S9**

**Tables S1 to S8**

**SI References**

Table of contents

Text S1. Geochemistry

Table S1

Table S2

*Sedimentary ancient DNA samples*

Table S3

Figure S1

Figure S2

Figure S3

Text S2. Fecal biomarker analysis

*Samples*

*Analytical protocol*

*Gas Chromatography-Mass Spectrometry*

Table S4

*Calculation of biomarker ratios*

*Concentrations of stanols and Δ^5^-sterols*

*Ratios of stanols and Δ^5^-sterols*

*Concentrations of bile acids*

*Discussion*

Figure S4

Figure S5

Figure S6

Text S3. Stable isotope analysis

*Application*

Table S5

Table S6

Text S4. Pollen analysis

*Laboratory work and analysis*

*Results and interpretation*

Table S7

*Low magnification analysis*

Table S8

*South Mesna lake as archive for pollen and its catchment area*

*Comments regarding Cannabis-type (Cannabinaceae) and Humulus*

Figure S7

Figure S8

Figure S9

Supplementary references

**Text S1. Geochemistry**

This section concerns the analysis of sediment core samples from Sør-Mesna (hereafter referred to as South Mesna lake) and Nord-Mesna (hereafter referred to as North Mesna lake) retrieved at the end of June 2020, as well as nine ^14^C AMS samples. All nine samples are from the core SMP320 retrieved from the southern basin of South Mesna lake. An overview of all the cores retrieved from both lakes can be found in table S1.

***Table S1.*** *An overview of the coordinates and depths (according to the* ***L****owest* ***R****egulated* ***W****ater level (LRW)) for cores retrieved from South- and North Mesna lake.* ***P*** *in the core name refers to long cores retrieved by a piston corer mounted to a pontoon raft, while* ***U*** *refers to cores retrieved from undisturbed surface sediments down to a depth of 1 meter with a UWITEC-corer. A note is also made whether one reached glacial deposits.*

| **Cores from South Mesna lake** | | | | |
| --- | --- | --- | --- | --- |
| **Core name** | **Date** | **Depth / under LRW** | **Coordinates** | **Notes** |
| SMP-120 | 16.06.2020 | 13.5 m / 5.5 m | 61.091657°N 10.706920°E | To deglaciation |
| SMU-120 | 16.06.2020 | 13.5 m / 5.5 m | 61.091657°N 10.706920°E | Surface sediments |
| SMP-220 | 16.06.2020 | 13.0 m / 5.0 m | 61.090329°N 10.709431°E | To deglaciation |
| SMU-220 | 16.06.2020 | 13.0 m / 5.0 m | 61.090329°N 10.709431°E | Surface sediments |
| SMP-320 | 17.06.2020 | 15.0 m / 7.0 m | 61.072324°N 10.819734°E | To deglaciation |
| SMU-320 | 17.06.2020 | 15.0 m / 7.0 m | 61.072324°N 10.819734°E | Surface sediments |
| SMP-420 | 17.06.2020 | 23.5 m / 15. 5 m | 61.079583°N 10.780789°E | **Not** to deglaciation |
| SMU-420 | 17.06.2020 | 23.5 m / 15.5 m | 61.079583°N 10.780789°E | Surface sediments |
| **Cores from North Mesna lake** | | | | |
| **Core name** | **Date** | **Depth / under LRW** | **Coordinates** |  |
| NMP-120 | 18.06.2020 | 29.0 m / 20.0 m | 61.104380°N 10.667056°E | To deglaciation |
| NMU-120 | 18.06.2020 | 29.0 m / 20.0 m | 61.104380°N 10.667056°E | Surface sediments |
| NMP-220 | 18.06.2020 | 14.5 m / 5.5 m | 61.101434°N 10.667540°E | To deglaciation |
| NMU-220 | 18.06.2020 | 14.5 m / 5.5 m | 61.101434°N 10.667540°E | Surface sediments |

In total, six piston cores and six gravity cores were marked, split, cleaned, described and analyzed. This work was performed by Joseph M. Buckby with the guidance of Professor Svein Olaf Dahl. Magnetic susceptibility (MS) has been measured, and photos were taken to effectively and reliably correlate between the different cores. It was decided with researchers from the Museum of Cultural History that the long core from the southern-most basin of South Mesna lake, SMP320, were to be selected for more comprehensive analyses related to element analysis using XRF (X-Ray Fluorescence), loss-on-ignition (LOI), dry-bulk-density (DBD) with the highest resolution sampling occurring using MS (Dataset S1). This work was, at times, heavily impacted by closed-down laboratories and the lack of (necessary) periodic maintenance of instruments due to the COVID-19 pandemic.

Figure S1 displays a graph with LOI and DBD from SMP320. The diagram shows an overall high amount of organic content only interrupted by short sequences of more minerogenic content. These episodes are primarily interpreted to have been caused by flood events in the rivers that flow into the southern end of South Mesna lake (northern origin). These events can be observed visually in the core as light layers against a background of darker more organic sediments. The LOI values are low and the DBD values are higher than the surrounding sediments (they mirror each other). Concurrently the MS values are high and additionally the values for titanium (Ti) and aluminum (Al) from the XRF-analysis are mostly positive in these “flood-layers”.

Some disturbances can be noted at the bottom of SMP320 in the transition between glacial deposits to Holocene sediments with a high organic content. This can be due to landslides in the basin and can possibly be correlated to a possible earthquake in the area due to the deglaciation in the Lillehammer region. As these sediments consist of easily deformable clayey silt, the disturbances may also be the result of the sampling process.

Obtaining age control was extremely important for this project and a consensus was reached to concentrate all radiocarbon samples to the main core, i.e. SMP320. There is an abundance of plant macrofossils in the cores of both lakes, but a closer examination revealed that most of this material was of aquatic origin. This material is less reliable for ^14^C AMS dating due to the possibility of a freshwater reservoir effect. A considerable effort has therefore been made to find terrestrial plant macrofossils from the different layers. After a thorough examination enough terrestrial plant macrofossils were found in nine different layers in SMP320 (table S2). The radiocarbon dating was performed at Poznan Radiocarbon Laboratory in Poznan, Poland.

***Table S2.*** *The ^14^C AMS datings of the nine terrestrial plant macrofossils as well as their inferred 2-σ age ranges.*

| **Sample name** | **Depth** | **Lab. Nr.** | **Age (uncal BP)** | **Note** | **2-σ calendar age ranges (cal. BP)** |
| --- | --- | --- | --- | --- | --- |
| SMP-320 | 21-21.5 | Poz-138576 | 880±30 | 0.4mg C | 703 - 1275 |
| SMP-320 | 35-35.5 | Poz-138577 | 2285±30 | 0.7mg C | 2045 – 2379 |
| SMP-320 | 89-89.5 | Poz-138578 | 4160±35 |  | 4442 – 4808 |
| SMP-320 | 114-114.5 | Poz-138579 | 4455±35 | 0.7mg C | 4916 – 5271 |
| SMP-320 | 135-135.5 | Poz-138581 | 4700±35 |  | 5324 – 5568 |
| SMP-320 | 155-155.5 | Poz-138582 | 4840±40 |  | 5547 - 5940 |
| SMP-320 | 178-178.5 | Poz-138583 | 6050±40 |  | 6688 – 7027 |
| SMP-320 | 230.5-231 | Poz-138584 | 7510±40 |  | 8165 – 8431 |
| SMP-320 | 250.5-251 | Poz-138585 | 8150±40 |  | 8682 - 9227 |

*Sedimentary ancient DNA samples*

In addition to high-resolution sediment analyses of SMP320 are also three freshwater sediment samples collected for aDNA analysis, and 32 samples chosen for pollen analysis. The aDNA samples were collected from a depth of 27, 115 and 267cm in dialogue with the Museum of Cultural History (Fig. S2), while the pollen samples were retrieved from 0-1cm, and then from 10cm down to the bottom of the core (see text S4 for report on pollen analysis). The aDNA samples were examined for brown trout (*Salmo trutta)*. A trace small amount of DNA matching was identified for brown trout and the results were therefore not further utilized. For further elaboration contact the corresponding author. Processing of ancient DNA and data analysis were performed by the SciLifeLab Ancient DNA facility. The sequencing was performed by the SNP&SEQ Technology Platform in Uppsala, part of the National Genomics Infrastructure (NGI) Sweden and Science for Life Laboratory. The SNP&SEQ Platform is also supported by the Swedish Research Council and the Knut and Alice Wallenberg Foundation.

***Table S3.*** *A summation of the results of the three aDNA samples including the total number of merged DNA reads generated and the number of reads successfully mapped to the brown trout genome.*

| **Facility sample ID** | **User sample description** | **Total no. of merged DNA**  **reads generated** | **DNA reads mapped to**  **brown trout genome** |
| --- | --- | --- | --- |
| A049-001 | aDNA-a, SMP320/27cm | 29,334,589 | 3,331 |
| A049-002 | aDNA-b, SMP320/115cm | 32,363,294 | 2,737 |
| A049-003 | aDNA-c, SMP320/267cm | 33,888,243 | 4,675 |

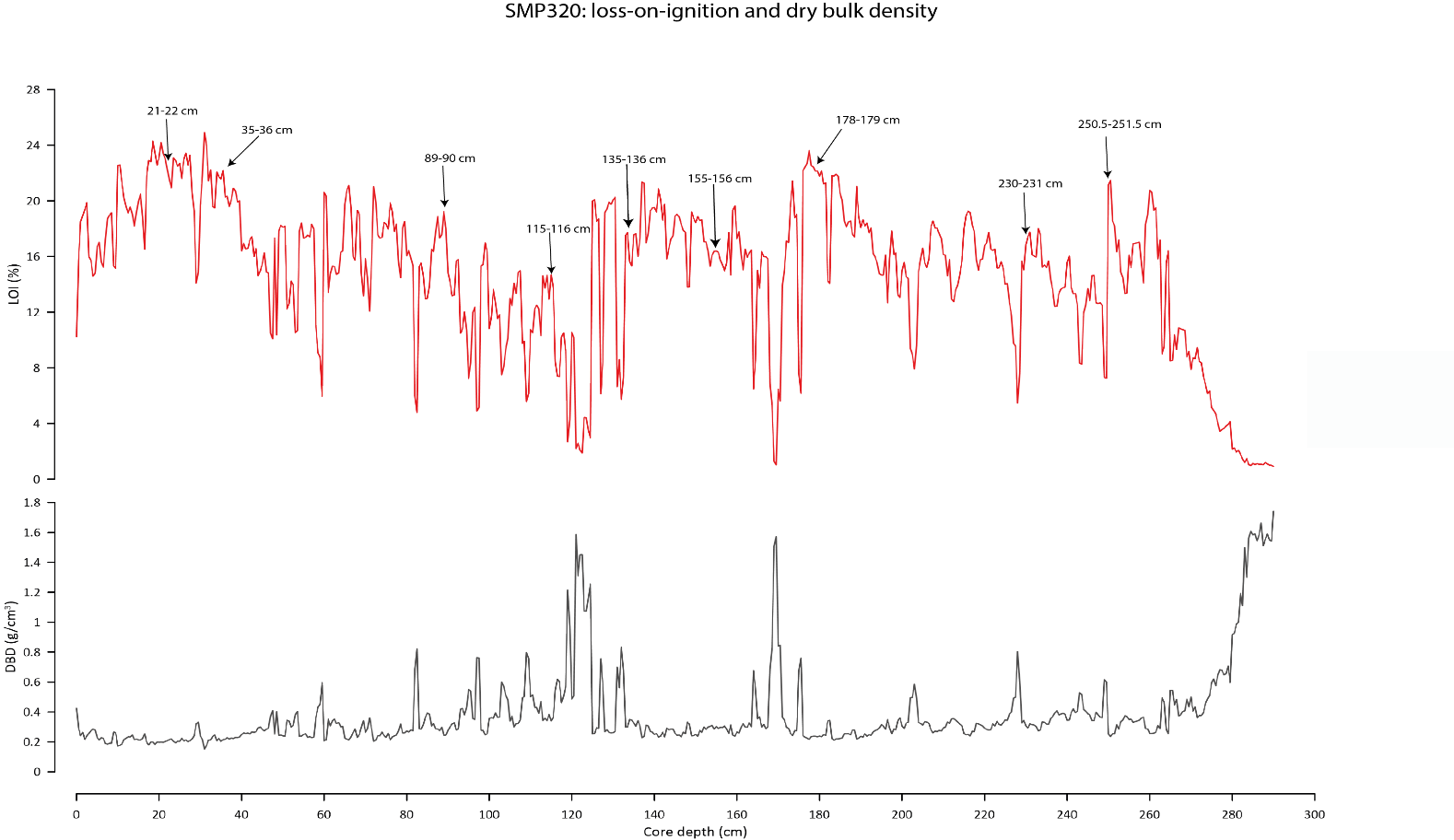

Figure S1. Loss-on-ignition (LOI) and dry-bulk-density (DBD) from deglaciation to early Holocene in core SMP-320 from the southern end of South Mesna lake. LOI and DBD are meant to “mirror” each other. The big fluctuations in LOI/DBD throughout most of the core are likely due to major flooding events. The bottom half of the core may be disturbed by a massive land slide. The nine ^14^C- samples are marked.

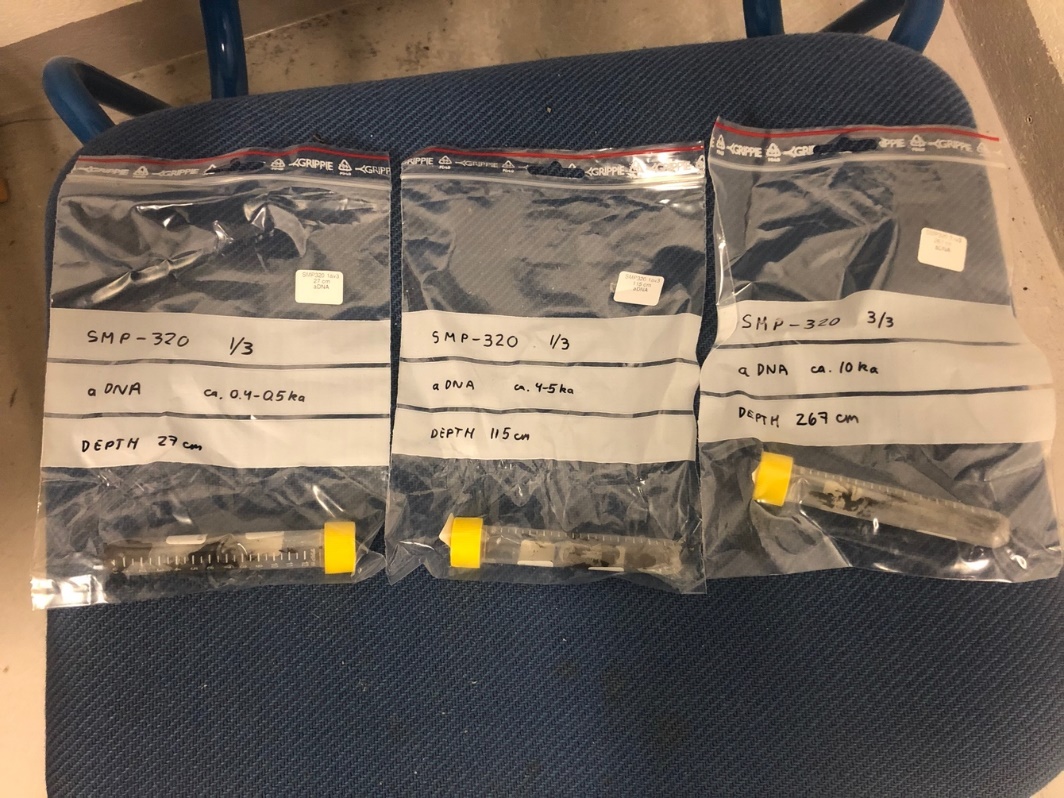

*Figure S2. aDNA samples from SMP-320. Estimated ages for the three samples are marked. Double samples from each layer were obtained to ensure that there was enough material for the analysis and strict rules for sampling were employed to prevent contamination of the material.*

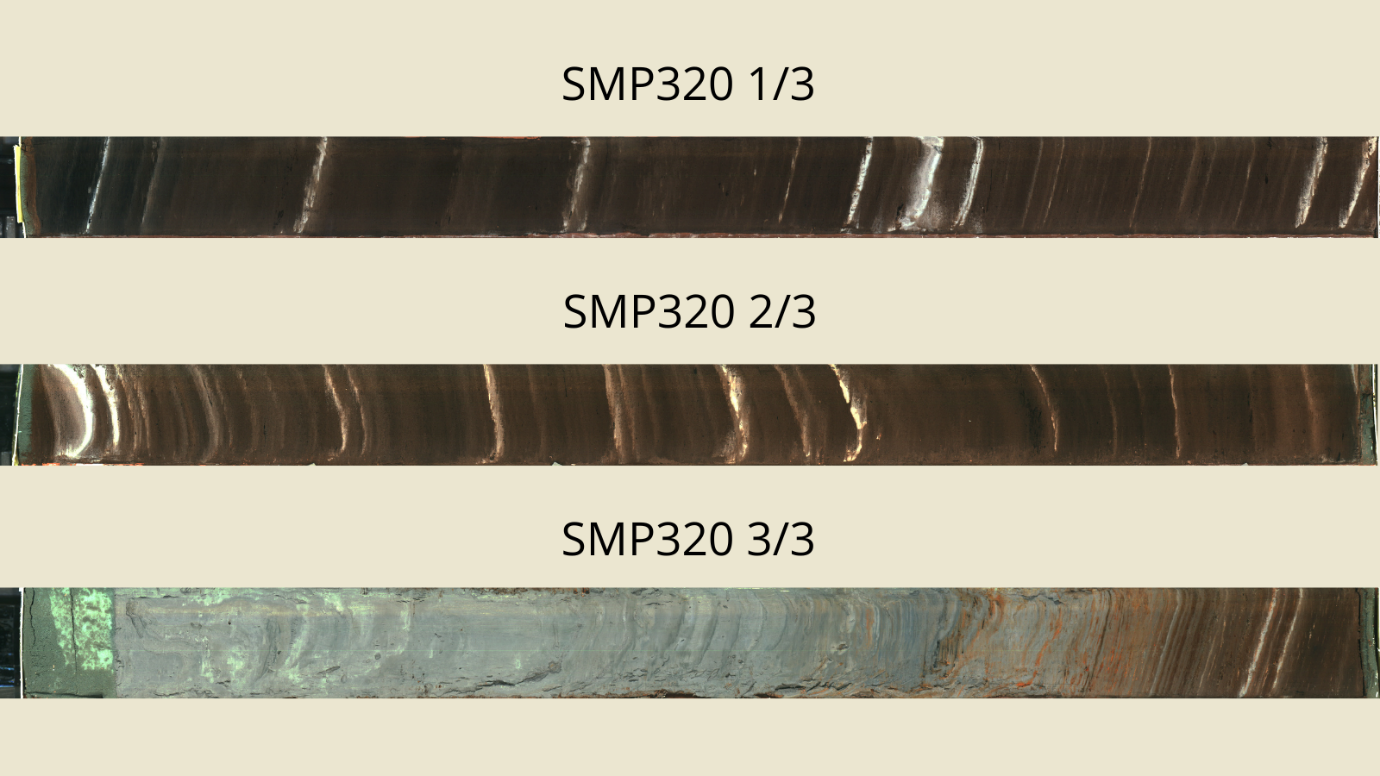

Figure S3. Photos of bright high-resolution sediment core sections, illustrating the visual variations across the core.

**Text S2. Fecal biomarker analysis**

Analyses of fecal stanols and bile acids can provide information about the presence of past human and animal populations when recovered from sedimentary deposits. Fecal stanols are organic molecules produced in the guts of mammals, most especially humans, who produce high amounts of coprostanol relative to other animals, and can be used to detect relative population levels on a landscape level (1-4). For example, fecal stanols have recently been used at the Mississippian site of Cahokia to corroborate population and land-clearance models associated with urbanism and mound construction (5) and the presence of Ice Age foragers in Beringia prior to the Last Glacial Maximum (6). The combination of fecal stanols and bile acids permits for the identification of different species of mammals and relative numbers of their presence, which are valuable in reconstructing land-use scenarios over time (7-9).

Specifically, 5β-stanols coprostanol and 5β-stigmastanol are fecal biomarkers produced in the guts of mammals from the membrane components of Δ^5^-sterols (e.g. cholesterol and β-sitosterol) (Figure S3). All fecal biomarkers are found in the feces of omnivores and in the feces of herbivores. However, there are differences between different animal taxa.

Coprostanol (a reduction product of cholesterol) is highly concentrated in the feces of omnivores (humans and pigs) in relation to other stanols 5β-stigmastanol (a reduction product of the plant membrane component β-sitosterol and further plant Δ^5^-sterols) is highly concentrated in the feces of herbivores in relation to other stanols. 5β-stanols can be further transformed in the gut and in the environment after deposition to their 3α-isomers (epi-coprostanol and epi-5β-stigmastanol). These substances are also biomarkers for feces.

Ratios of these biomarkers in relation to one another are used to track the presence of ruminants and monogastric animals within sedimentary contexts. In settings like Norway, monogastric animals can include feline and canine carnivores, pigs (*Suid* sp.) and humans. However, carnivores produce only small amounts of 5β-stanols and excrete Δ^5^-sterols [for](https://www.dict.cc/?s=for) [the](https://www.dict.cc/?s=the) [most](https://www.dict.cc/?s=most) [part](https://www.dict.cc/?s=part) untransformed. Taking transport and taphonomic concerns into account, generally speaking, the more molecules that are detected in the analysis, the higher the population of the proxy taxa can be inferred (10-12).

Bile acids are assayed to obtain information about different omnivorous feces. Hyodeoxycholic acid (HDCA) is the most common bile acid in pig feces (wild boar, in the context of pre-Iron Age Scandinavia). In the feces of humans, HDCA represents a negligible component of the bile acid pool (4, 9).

*Samples*

A total of seven samples from SMP320 were studied (Dataset S1) as well as a sample collected from modern beach mud at the landing dock of the research pontoon in 2020 (labelled MES20-3b). Samples were removed from the cores by David Wright at the Museum of Cultural History in Oslo in June and August 2020. They were then refrigerated prior to shipping to the Institute of Geography at the University of Mainz, Germany. A total lipid extraction was performed by microwave oven, followed by purification, derivatization and GC-MS following protocols outlined in Birk et al. (13).

*Analytical protocol*

For extraction the method of was Golinski (14) used and for the purification and measurement of stanols, Δ^5^-sterols and bile acids the method from Birk *et al.* (13) was used with small modifications. Standards of stanols, Δ^5^-sterols and bile acids and suppliers are listed in dataset S1. Cholesterol-d7 and lithocholic acid-d4 were obtained from CDN Isotopes (Pointe-Claire, Canada). 5α-Cholestan-3β-ol-d5, 5α-cholestane-d4 and derivatization reagents were obtained from Sigma-Aldrich (St. Louis, MO, USA). Distilled water was used, and all other chemicals were GC- or at least analytical grade and were purchased from various suppliers.

To extract the biomarkers by microwave-assisted extraction (Multiwave PRO, Anton Paar, Graz, Austria) 5 – 10 g finely grounded sediment was weighted into 100 ml PTFE-TFM liners. 40 mL dichloromethane/methanol (2:1; v/v) and a stirring bar were added, and the liners were placed in PEEK-GF vessels (MF100, Anton Paar, Graz, Austria) into the rotor (16SOLV, Anton Paar, Graz, Austria). Extraction was carried out with stirring. The temperature program was heated to 100 °C within 10 min, and held at 100 °C for 20 min. After cooling at room temperature, the suspension was transferred to a 70 mL centrifuge tube. Centrifugation was done at 3000 rpm for 5 min and the residue was washed two times with 40 ml dichloromethane/methanol each.

After extraction, deuterated cholestane, cholesterol and lithocholic acid were added as first internal standards (IS1). The extract was concentrated by rotary evaporation and completely dried under a gentle stream of nitrogen. Drying under a gentle stream of nitrogen was always done to dry extracts and is not repeatedly described in the following sections of the method description.

The extract was saponified in 5 ml 0.7 M KOH in methanol and reaction was allowed overnight (~15 h) at room temperature. After saponification, 10 ml water was added to the extract and the neutral fraction including the stanols and Δ^5^-sterols was extracted by repeated liquid–liquid extraction with chloroform (3 × 15 mL). To extract the bile acids, 6 M HCl was added to the saponification/water solution for acidification (pH ≤2) and the acidic fraction including the bile acids was extracted by repeated liquid–liquid extraction with chloroform (3 × 15mL). The chloroform extracts were then concentrated by rotary-evaporation and dried.

The acidic fraction was methylated in 1mL dry 1.25 M HCl in methanol at 80 °C for 2h. The methyl esters were extracted after addition of 1mL water by repeated liquid–liquid extraction with hexane (3 × 1.5ml), followed by drying of the extract.

Both the neutral and acid fractions were purified by solid phase extraction (SPE). Purification of the neutral fraction was carried out with columns (5mm diameter, PE) packed with 5% deactivated silica gel (50mm, pore size 100 Å, particle size 63–200µm, type POLYGOPREP 100-130, Macherey-Nagel, Düren, Germany) and preconditioned with hexane. Less polar substances including *n-*alkanes were eluted with 4mL hexane and dried. The fraction containing alcohols including stanols and Δ^5^-sterols, was eluted first with 3 mL dichloromethane and afterwards with 2 mL dichloromethane/acetone (2:1, v/v). The dichloromethane and dichloromethane/acetone eluates were combined and dried.

Purification of the acid fraction was carried out with columns (5mm diameter, PE) packed with activated silica gel (50mm, pore size 100 Å, particle size 63–200µm, type POLYGOPREP 100-130, Macherey-Nagel, Düren, Germany) and preconditioned with dichloromethane/hexane (2:1, v/v). Less polar substances (fatty acid methyl esters, etc.) were eluted with 4mL dichloromethane/hexane (2:1, v/v). The hydroxy acids including bile acids were eluted with 5mL dichloromethane/methanol (2:1, v/v) and the eluate was dried.

For silylation of stanols and Δ^5^-sterols, 37.5µL BSTFA+TMCS (99:1 v/v) and 12.5µL pyridine were added and the mixture was heated at 90°C for 1h. Excess silylation reagent was evaporated and dry toluene and 5α-cholestane (dissolved in dry toluene) as second internal standard (IS2) were added.

The fraction containing bile acids was silylated in 50 µL dry toluene, 98 µL BSTFA and 2 µL TSIM. After heating at 80 °C for 1 h, 5α-cholestane (dissolved in dry toluene) was added as IS2. 5α-cholestane (dissolved in dry hexane) was added as IS2 to the *n-*alkane fraction before measurement. External standard solutions of the relevant biomarkers and IS1 were prepared in methanol or hexane and derivatized using the same methods, which were used for the soil extracts.

*Gas Chromatography-Mass Spectrometry*

Derivates of stanols, Δ^5^-sterols and bile acids were analyzed via gas chromatography-mass spectrometry (GC/MS) with an Agilent 7000D mass spectrometer (Agilent, Santa Clara, CA, USA) connected to an Agilent 7890B gas chromatograph (Agilent, Santa Clara, CA, USA). A DB-5ms Ultra Inert (Agilent, Santa Clara, CA, USA), 30 m fused silica column with 0.25 mm I.D. and 0.25 µm film thickness was directly coupled with the mass spectrometer. The injection port was equipped with a split/splitless liner (5183-4711, Agilent, Santa Clara, CA, USA). He (99.9999% purity) was used as carrier gas at a column flow of 1 mL min*-*^1^ (constant flow). The transfer line temperature was 250 °C. Electron impact ionization was used at 70 eV. Measurements in scan mode were done to verify peak identity and measurements in selected ion monitoring mode (SIM) were performed for quantification (Table S4 shows the selected ions).

***Table S4.*** *Investigated compounds, suppliers, retention times (RT) and characteristic ion fragments.*

| Biomarker group | Substance | RT (min) | Characteristic ion fragments (m/z) |
| --- | --- | --- | --- |
| Stanols | 5β-cholestan-3β-ol^a^ | 26.2 | 370; 215; 355 |
|  | 5β-cholestan-3α-ol^b^ | 26.7 | 355; 215; 370 |
|  | 5α-cholestan-3β-ol^c^ | 28.5 | 445; 355; 460 |
|  | 5β-stigmastan-3β-ol^d^ | 30.8 | 398; 215; 383 |
|  | 5β-stigmastan-3α-ol^d^ | 31.5 | 398; 215; 383 |
|  | 5α-stigmastan-3β-ol^b^ | 33.9 | 215; 383; 398 |
| Δ^5^-sterols | cholest-5-en-3β-ol^b^ | 28.3 | 329; 368; 458  394; 255; 484  396; 357; 486 |
|  | stigmasta-5,22-dien-3β-ol^a^ | 31.7 |  |
|  | stigmast-5-en-3β-ol^b^ | 33.5 |  |
| Bile acids | 3β-hydroxy-5β-cholanoic acid (ILCA)^a^ | 26.7 | 215; 257; 357 |
|  | 3α-hydroxy-5β-cholanoic acid (LCA)^b^ | 27.6 | 215; 257; 372 |
|  | 3β,12α-dihydroxy-5β-cholanoic acid (BDCA)^a^ | 27.9 | 255; 345 |
|  | 3α,12α-dihydroxy-5β-cholanoic acid (DCA)^b^ | 28.4 | 255; 345; 370 |
|  | 3α,7α -dihydroxy-5β-cholanoic acid (CDCA)^b^ | 29.4 | 255; 355; 370 |
|  | 3α,6α-dihydroxy-5β-cholanoic acid (HDCA)^a^ | 30.0 | 255; 355; 370 |
|  | 3α,7β-dihydroxy-5β-cholanoic acid (UDCA)^b^ | 31.1 | 255; 370; 460 |

^a^ obtained from Steraloids (Newport, RI, USA)

^b^ obtained from Sigma-Aldrich (St. Louis, MO, USA)

^c^ obtained from Alfa Aesar (Ward Hill, MA USA)

^d^ obtained from Chiron (Trondheim, Norway)

For analyses of derivatives of stanols and Δ^5^-sterols, the injection port was set to 250°C and 1µL was injected in pulsed splitless mode (pulse pressure: 180kPa, pulse time: 1.01min). The column temperature program was 80°C (held 1.5min) to 260°C at 12°Cmin*-*^1^, to 271°C at 0.5°Cmin*-*^1^ and to 300°C (held 10min) at 10°Cmin*-*^1^. For analyses of bile acid derivatives, the injection port was set to 290°C and 1µL was injected in splitless mode. The column temperature program was 80°C (held 1.5 min) to 250 °C at 20 °C min*-*^1^, to 280°C at 1.2°Cmin*-*^1^ and to 300°C (held 10min) at 10°Cmin*-*^1^. In case of high concentrations of biomarker derivatives, additional measurements were conducted in pulsed split mode and split mode, respectively.

*Calculation of biomarker ratios*

Cholesterol and plant-derived Δ^5^-sterols are mainly reduced to 5α-stanols (cholestanol and 5α-stigmastanol) in the environment, and a small part is reduced to 5β-stanols. If the results show that there were background values of coprostanol and other 5β-stanols in the samples from the feces of wild animals and other plant-based derivative processes a background correction is made. For South Mesna lake, to try to correct for background values, ratios for omnivorous animals (pigs, humans) were calculated based on the following two equations:

“omnivore1” = coprostanol/cholestanol

“omnivore2” = (coprostanol + epi-coprostanol)/cholestanol

The numerators of these equations are biomarkers of feces, whereas the denominators are the molecules reduced in the sediments. High ratios indicate whether there was an input of omnivorous feces.

A similar ratio was calculated with the fecal biomarker that is highly concentrated in the feces of herbivores (ruminants) with the equation:

“herbivore1” = 5β-stigmastanol/5α-stigmastanol

A ratio including the epi-form of the fecal biomarker that is highly concentrated in the feces of herbivores (cattle, goat, sheep) was not calculated because this substance was detected only in a few samples.

The ratio omnivore/herbivore1 was calculated to differentiate between fecal inputs from omnivores and herbivores. To distinguish between the two, the quantity of fecal biomarkers that are highly concentrated in the feces of omnivores was divided by the quantity of fecal biomarkers that are highly concentrated in the feces of herbivores.

“omnivore/herbivore1” = coprostanol/5β-stigmastanol

The numerator of this equation is the biomarker of omnivore feces, whereas the denominator is the biomarker of herbivore feces. This ratio is only useful if fecal matter is present in the samples and fecal biomarkers are above the background values. Concentrations of fecal biomarkers in samples are reported in relation to total organic carbon (TOC) and to dry weight of the sediment.

*Concentrations of stanols and Δ^5^-sterols*

The amounts related to TOC of the fecal biomarker coprostanol, which is found in high concentrations in human feces, were ≤13µg g_TOC_^-1^ and the biomarker that is characteristic for herbivore feces, 5β-stigmastanol, had concentrations in TOC ≤34µg g_TOC_^-1^ (Fig. S4; A, B). The 3α-epimers of coprostanol and 5β-stigmastanol (epi-coprostanol and epi-5β-stigmastanol) were not found in all samples and had amounts in TOC ≤10µg g_TOC_^-1^ (Fig. S4; C, D).

Highest concentrations of fecal biomarkers were found in sample 1/3-10 (125cm). Concentrations of coprostanol, epi-coprostanol and 5β-stigmastanol equal to or above the median were found in addition in the samples 2/3-4 (174.75cm) and 3/3-5 (333.5cm) (Fig. S4; A – C).

The steroids, which originate from animal and plant membranes, were found to have significantly elevated concentrations as fecal biomarkers. The amounts of cholesterol and stigmasterol related to TOC were ≤69µg g_TOC_^-1^ and β-sitosterol had concentrations in TOC ≤322 µg g_TOC_^-1^. (Fig. S4; G – I). The substances that were built from these compounds by reduction processes in the environment (outside of mammal guts) had also higher concentrations than the fecal biomarkers. The concentrations in TOC of cholestanol were ≤42µg g_TOC_^-1^ and of stigmastanol were ≤140µg g_TOC_^-1^ (Fig. S4; E, F). In contrast to the fecal steroids, the individual 5α-stanols and Δ^5^-sterols had their highest concentrations in different samples and did not show a common maximum in a specific depth (Fig. S4; E – I).

*Ratios of stanols and Δ^5^-sterols*

To correct for background levels of stanols, ratio values were calculated. The ratios, that are enhanced due to feces input “omnivore1” (coprostanol/cholestanol), “omnivore2” ((coprostanol + epi-coprostanol)/cholestanol) and “herbivore1” (5β-stigmastanol/5α-stigmastanol) had values ≤53 (Fig. S5; A – C). As with the concentrations of fecal biomarkers, these ratios were found to be equal to or above the median in samples 1/3-10 (125cm), 2/3-4 (174.75cm) and 3/3-5 (333.5cm) and the highest values were found in sample 1/3-10 (125cm) (Fig. S5; A – C).

The ratio to differentiate between feces of omnivores and herbivores “omnivore/herbivore1” (coprostanol/5β-stigmastanol) was ≤0.50 (Fig. S5; D). The median of ratio “omnivore/herbivore1” was 0.23 and both samples with enhanced concentrations of fecal biomarkers and biomarker ratios in the middle of the core (1/3-10 (125cm), 2/3-4 (174.75cm)) had values that were above the median (0.38 in sample 1/3-10) or only slightly below the median (0.22 in sample 2/3-4) (Fig. S5; D). However, the lowest sample 3/3-10 had a low value of 0.14 (Fig. S5; D).

*Concentrations of bile acids*

Bile acids yielded concentrations at background levels in the middle and lower part of the core, although the characteristic biomarker of pigs (hyodeoxycholic acid) was not identified at all (data not shown).

*Discussion*

The concentration patterns of stanols and Δ^5^-sterols in the samples, which showed plant-membrane derived steroids (β-sitosterol, stigmasterol and stigmastanol) in high concentrations, middle concentrations of cholesterol and cholestanol and low concentrations of fecal steroids are typical for soils (13). The amounts of fecal stanols were comparable to the concentration reported by d’Anjou et al. (15) from a lake in northern Norway (≤20μg g_TOC_^-1^).

The enhanced concentrations of fecal biomarkers combined with enhanced ratios that show fecal inputs in the samples 1/3-10 (125cm), 2/3-4 (174.75cm) and 3/3-5 (333.5cm) indicate higher input of feces in these samples in comparison to the other samples. Both the ratios indicative of omnivorous feces and the ratios indicative of herbivorous feces were elevated in these samples. Since the biomarkers that dominate omnivore feces are also present in herbivore feces and vice versa, these results are not contradictory (Fig. S3). It is also possible that both types of feces were entered. However, the ratio to differentiate between omnivore and herbivore feces showed that possible fecal inputs in the middle part of the core had a higher proportion of omnivore feces than in the lowest sample, which had a comparatively high proportion of herbivore feces.

The low concentrations of bile acids do not allow detailed interpretation because the patterns are typical for an unspecific background (JJB, own unpublished data). The typical bile acid for the feces of pigs, HDCA, shows a low sensitivity in GC/MS measurements in comparison to the other bile acids (JJB, own unpublished data). However, in pig feces this bile acid occurs in high concentrations and the concentration of this bile acid is considerable higher than the concentration of coprostanol in pig feces (Fig. S3). Therefore, the absence of HDCA in the measurements could be a weak indication that the potential omnivore fecal inputs in the middle part of the core could originate from humans.

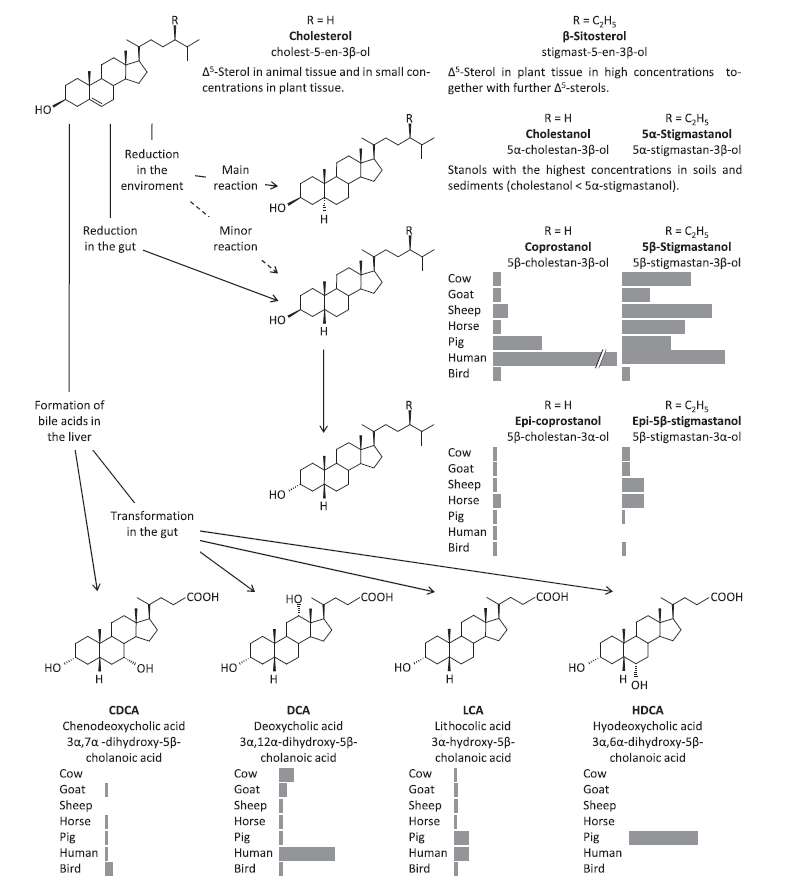

Figure S4. Formation of fecal biomarkers 5β-stanols (coprostanol, 5β-stigmastanol, epi-coprostanol and epi-5β-stigmastanol), bile acids and 5α-stanols following digestion and decomposition in sediments of Δ5-sterols (7).

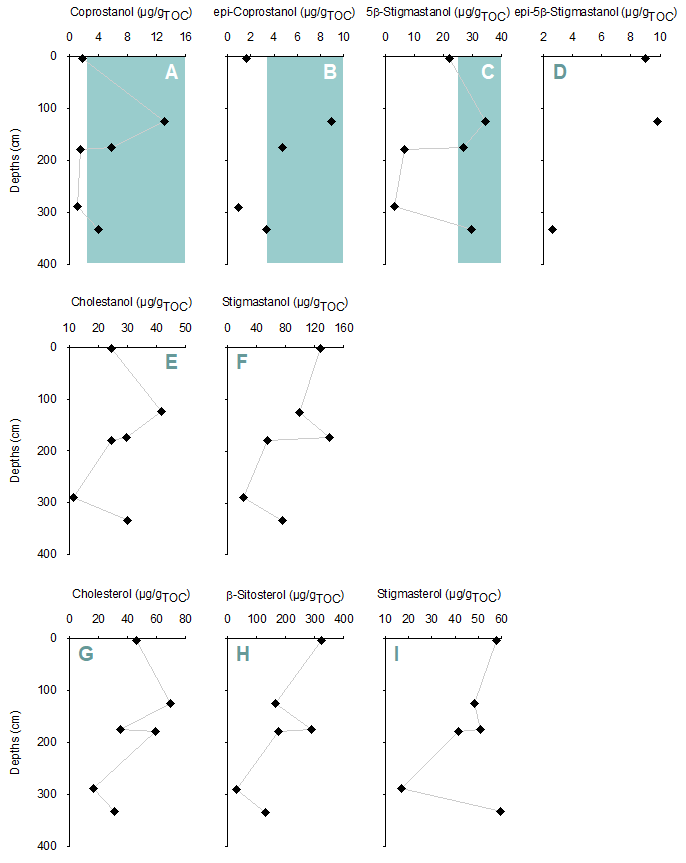

Figure S5. Concentrations related to TOC of, A, B, C, D, fecal stanols, E, F, stanols that are mainly build in the environment and, G, H, I, Δ^5^-sterols based on their places within SMP320 core.

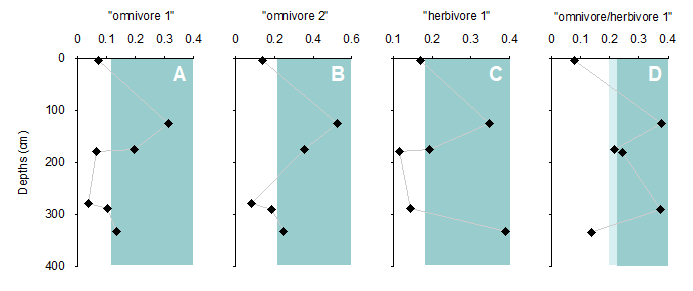

Figure S6. Calculated ratios of, A, B, gut-derived stanols from omnivorous animals divided by the corresponding environmental stanol, C, of a gut-derived stanol from herbivorous animals divided by the corresponding environmental stanol and, D, of gut-derived stanols from omnivorous vs. herbivorous animals from SMP320 core.

**Text S3. Stable isotope analysis**

All samples for study of stable carbon ($\delta$^13^C) and nitrogen ($\delta$^15^N) isotopes were normalized using an agate mortar and pestle. Sediments destined for analysis of $\delta$^13^C were saturated in 1M HCl for 24-48 hours on a shaker table to remove carbonates. All samples were visually inspected for the presence of bubbles to ensure complete removal of carbonates, and none of the samples were deemed to have remaining carbonates. Following the HCl treatment, the samples were rinsed, centrifuged and decanted three times in distilled H_2_O and dried in an oven at 40ºC for three days to ensure that they were thoroughly dried. Samples for $\delta$^15^N were not acidified. All samples were weighed on a microbalance to 25±1mg and packed into sterile tin capsules, with weights recorded to ±0.01mg. The samples were then taken to the CLimate Interpretation of Plant Tissue (CLIPT) laboratory in the Department of Biosciences at the University of Oslo.

Soil Organic Matter (SOM) was analyzed using a Flash 1110 Elemental Analyzer (EA) with a no-blank autosampler connected to a Delta V plus isotope ratio mass spectrometer. Samples were combusted with excess of O_2_ at 1020ºC, catalyzed with chromium oxide and silvered cobaltous oxide. N-oxides were reduced with reduced copper at 650ºC. A CO_2_ trap was used for the $\delta$^15^N analyses, and a water trap for both elements. Sodium percholate was used to clean the EA between runs of $\delta$^13^C and $\delta$^15^N to ensure no contamination of the instrument from CO “cracking” during $\delta$^13^C analysis, which can result in residual mass 28 molecules (^12^C + ^16^O) that could appear as ^14^N_2_ during $\delta$^15^N analysis. One set of triplicates was analyzed for every 10 samples, determined randomly. Standards used were IAEA-N1 and IAEA-N2 for N isotopes and USGS-24, IAEA-600, and IAEA-CH-6 for C isotopes (<https://analytical-reference-materials.iaea.org/>). Precision of standards were <0.2 per mille (‰) (table S5).

**Table S5*.*** *δ^13^C_VPDB_ internal references and quality assurance calibrated against NBS19 and LSVEC with consensus values 1.95‰ and -46.6‰, respectively. δ^15^N_AIR_ internal references and quality assurance calibrated against USGS40 and USGS41 with consensus values -4.52‰ and 47.57‰, respectively*.

| Internal reference and quality assurance material | Name | δ^13^C_VPDB_ Calibrated Value (‰) | δ^13^C_VPDB_ Measured Value (‰) | δ^13^C_VPDB_ Stdev | δ^13^C_VPDB_ n |
| --- | --- | --- | --- | --- | --- |
| Reference 1 | JGLUT | -13.43 |  |  |  |
| Reference 2 | POPPGLY | -36.58 |  |  |  |
| Quality assurance | JALA | -20.62 | -20.57 | 0.05 | 8 |
| Internal reference and quality assurance material | **Name** | **δ^15^N_AIR_ Calibrated Value (‰)** | **δ^15^N_AIR_ Measured Value (‰)** | **δ^15^N_AIR_ Stdev** | **δ^15^N_AIR_ n** |
| Reference 1 | JGLUT | -4.43 |  |  |  |
| Reference 2 | POPPGLY | 11.25 |  |  |  |
| Quality assurance | JALA | -3.16 | -3.26 | 0.04 | 7 |

*Application*

The results of stable isotope analysis of carbon and nitrogen (table S6) form the basis of interpreting formation and organic source material present on the landscape. There are two primary photosynthetic pathways that plants convert sunlight into sugars: one involves the production of a 3-carbon acid that is formed after splitting CO_2_ during carboxylization to produce the glucose that meets a plant’s metabolic requirements. The second pathway produces a 4-carbon glucose molecule through the Hatch-Slack photosynthetic process. Biochemically, these two processes result in differential conservation of ^13^C, with C3 plants (mostly trees) discriminating against ^13^C more than C4 plants (mostly grasses). Since Scandinavia is generally depauperate of C4 vegetation (16), enriched/depleted ^13^C values from soils and sediments that hosted terrestrial vegetation are generally interpreted as reflecting open/closed forest conditions, respectively, vis-à-vis the ‘canopy effect’ (17, 18). The canopy effect occurs when ^13^C-depleted CO_2_ is recirculated within the understory of closed forests and the attenuation of light changes photosynthetic activity and stomatal conductance of CO_2_ (19, 20). Thus, all other considerations held equal, the isotopic effect on soils that host vegetation is relatively linear in that grassier, more open conditions tend to have enriched (less negative) $\delta$^13^C values relative to more forested, closed conditions, which tend to be more depleted (more negative).

On the other hand, the depletion of the heavier nitrogen isotope (^15^N) in soils and sediments occurs when organic matter decomposition rates are high and ^15^N-depleted NH_4_^+^ is formed (21). During nitrification—a process in which ammonia (NH_3_) is aerobically converted to nitrite (NO_2_^-^)—concentrations of ^15^N increase in soils due to fractionation (22). Clear cutting of forests tend to demonstrate an initial increase in $\delta$^15^N accumulation in O-horizons of soils, however within 15 years following deforestation, $\delta$^15^N is the same as pre-clearance levels (21, 23). On the other hand, landscape burning, foddering and grazing of livestock tend to increase $\delta$^15^N, at least temporarily (24), but studies of anthropogenic soils from within pastoral encampments in Kenya have demonstrated long-term enrichment of $\delta$^15^N even in abandoned areas (25, 26). Similar effects have been detected in northern European plaggen soils (plaggosols), which are manured agricultural fields (27). Higher $\delta$^15^N in soils are also correlated with more arid surface and plant foliar conditions (28) as well as warmer ambient air temperatures (29).

**Table S6.** Results of the stable isotope analysis at the CLIPT laboratory, University of Oslo.

| **ID** | **Depth** | **δ^13^C_VPDB_ (‰)** | **%C** | **δ^15^N_AIR_ (‰)** | **%N** | **2-σ calendar age ranges (cal. BP)** |
| --- | --- | --- | --- | --- | --- | --- |
| 1/3-1 | 3cm | -28.38 | 13.23 | 2.20 | 0.71 | 178 – 923 |
| 1/3-2 | 19cm | -28.95 | 3.65 | 2.24 | 0.45 | 585 – 1199 |
| 1/3-3 | 32cm | -28.85 | 4.64 | 2.23 | 0.78 | 1493 – 2225 |
| 1/3-4 | 51cm | -28.58 | 11.48 | 2.27 | 0.74 | 2524 – 3603 |
| 1/3-5 | 65cm | -28.71 | 13.95 | 2.27 | 0.75 | 2950 – 4123 |
| 1/3-6 | 78cm | -29.33 | 12.53 | 2.41 | 0.73 | 3717 – 4538 |
| 1/3-7 | 94cm | -28.96 | 12.38 | 1.54 | 0.59 | 4529 – 4910 |
| 1/3-8 | 109cm | -28.68 | 2.75 | 1.86 | 0.50 | 4797 – 5189 |
| 1/3-9 | 113cm | -28.62 | 8.97 | 1.50 | 0.45 | 4902 – 5241 |
| 1/3-10a | 125cm | -27.83 | 0.52 | 1.04 | 0.11 | 5088 – 5446 |
| 1/3-10b | 125cm | -27.76 | 0.55 | 0.81 | 0.10 | 5088 – 5446 |
| 1/3-10c | 125cm | -27.77 | 0.54 | 0.77 | 0.10 | 5088 – 5446 |
| 2/3-1 | 133cm | -29.19 | 14.04 | 2.68 | 0.76 | 5282 – 5538 |
| 2/3-2 | 147cm | -29.32 | 12.46 | 2.60 | 0.73 | 5437 – 5800 |
| 2/3-3 | 172cm | -29.17 | 11.85 | 2.65 | 0.73 | 6281 – 6843 |
| 2/3-4 | 175cm | -27.30 | 0.20 | 0.43 | 0.04 | 6465 – 6939 |
| 2/3-5 | 180cm | -29.05 | 13.29 | 2.65 | 0.83 | 6742 – 7149 |
| 2/3-6 | 190cm | -29.80 | 14.29 | 2.59 | 0.78 | 6921 – 7542 |
| 2/3-7 | 221cm | -29.74 | 14.50 | 2.05 | 0.77 | 7735 – 8270 |
| 2/3-8 | 241cm | -30.11 | 7.74 | 1.60 | 0.67 | 8409 – 8939 |
| 2/3-9a | 257cm | -30.40 | 13.49 | 0.83 | 0.91 | 8795 – 9421 |
| 2/3-9b | 257cm | -30.39 | 12.29 | 0.85 | 0.94 | 8795 – 9421 |
| 2/3-9c | 257cm | -30.42 | 11.32 | 1.21 | 0.95 | 8795 – 9421 |
| 3/3-1a | 266cm | -30.59 | 17.44 | 0.95 | 0.85 | 8945 – 9723 |
| 3/3-1b | 266cm | -30.52 | 8.98 | 1.01 | 0.83 | 8945 – 9723 |
| 3/3-1c | 266cm | -30.41 | 8.70 | 1.18 | 0.80 | 8945 – 9723 |
| 3/3-2 | 279cm | -28.98 | 3.63 | 1.55 | 0.36 | 9151 – 10,079 |
| 3/3-3 | 290cm | -26.89 | 0.22 | -0.05 | 0.04 | 9335 – 10,367 |
| 3/3-4 | 297cm | -26.89 | 0.17 | -0.80 | 0.04 | 9456 – 10,546 |
| 3/3-5 | 334cm | -26.06 | 0.21 | -0.48 | 0.07 | 10,064 – 11472 |
| 3/3-6 | 369cm | -25.68 | 0.24 | -0.86 | 0.04 | 10,684 – 12,348 |

**Text S4. Pollen analysis**

Pollen analyses were conducted on samples from one sediment core (SMP320) from the southern end of South Mesna lake in Ringsaker municipality. The intention of the analyses was to investigate the vegetation development and the land use in the area from deglaciation to present. Specific questions were raised concerning changes in the proxy data under the early Neolithic (approx. 5500 – 4700 cal. BP, 110-150cm deep) that could be linked to human activity and land use.

The locality is a large lake that is situated in a valley just over 520 m.a.s.l. with a big part of the drainage area, thus also the pollen uptake area, up to 100m higher than the lake surface. The lake is over 7km long in the NW-SE direction, approx. 1km wide, and in places have well-developed sandy-gravel beach zones due to the relatively high wave energy. The outlet is to the west through the Bustokk River. The water depth at sampling points SMP 320, in the SE part of the lake, was approx. 15m. Multiple smaller water courses drain into the eastern part of the lake from the northern, eastern, and southern directions. Multiple big wetlands are also located around the water course. Directly west of the Mesna lakes is the city of Lillehammer in Gubrandsdalen with Lake Mjøsa directly south. Lake Mjøsa is considerably lower at 122 m.a.s.l. and has today some elements of broadleaf forest in its immediate surroundings.

*Laboratory work and analysis*

The sediment samples taken out for pollen analysis were approximately 1cm^3^ and a total of 15 samples were analyzed from approximately every 20cm, from the depth range of 230cm to 10cm from the sediment core. The enrichment of the pollen samples was done in the microfossil laboratory at Arkeologerna, Stockholm, according to standard methods described by Berglund & Ralska-Jasiewiczowa (30). This essentially involves dispersion in sodium hydroxide (NaOH), removal of any calcium carbonate (CaCO_3_) with hydrochloric acid (HCl), removal of cellulose with an acetolysis solution consisting of concentrated sulfuric acid (H_2_SO_4_) and acetic anhydride (CH_3_COO)_2_O. When preparing preparations for microscopy, the samples were embedded in glycerin.

The identification of pollen and spores has been done with the help of reference literature (31). A reference collection of pollen preparations has been used when necessary. Pollen from land plants has been used as the pollen sum in the percentage calculation, i.e. trees, shrubs, dwarf shrubs and herbs. Ferns, water and marsh plants, as well as mosses and algae have been counted outside this total. In parallel with the pollen analysis, charcoal particles >20µm have also been counted and are presented as percentages of the pollen sum. The samples have been supplemented with a "low magnification analysis" to be able to detect the presence of low-frequency pollen types with generally large pollen grains, at all sample levels. The process means that a larger part of each sample is screened (in practice the rest of the microscope preparations) at lower magnification (100 x), but only large pollen grains such as grains (barley, rye, wheat and oats) are identified and counted.

The low-magnification analysis is very time-efficient and provides more continuous data of the affected taxa, and the possibility of detecting very low concentrations of pollen.

*Results and interpretation*

A total of 48 different pollen types (pollen taxa) from land plants were found during the analysis, and if pollen from aquatic plants and spores from vascular cryptogams are included, there were a total of 59 taxa. The pollen sums amounted to between 700–800 grains, except in the bottom sample where the pollen concentration was too low. The full analysis results are presented in table S7. Charcoal particles are presented in the table, but they were generally present in low concentrations but fortunately with some variation. Also, some algae such as *Pediastrum* sp. have been included in the table. Spores from bryophytes (mosses), mainly bear mosses (*Polytrichum* sp.) were abundant in all samples but were not counted. Indeterminate pollen, i.e. pollen grains that are mainly unidentifiable because they are corroded or deformed, are also included as a percentage in table S7. The proportion of indeterminate pollen is relatively high, between 4 and 10%, which probably reflects the erosion and re-deposition that occur in South Mesna lake’s beach zones, for example in connection with storm events. It is also likely that erosion and redeposition have taken place along the waterways that lead to the lake, for example peatlands may have been eroded in connection with spring floods or events with high flows. The most important pollen types for interpretation of the local vegetation and landscaping are presented in a pollen diagram (Figure S6). Charcoal has also been included in the diagram; however, the levels of charcoal are generally very low in the layer sequence.

| **epth (cm)** |  | 10 | 30 | 50 | 70 | 90 | 110 | 120 | 140 | 160 | 180 | 200 | 230 | 250 | 270 | 290 |
| --- | --- | --- | --- | --- | --- | --- | --- | --- | --- | --- | --- | --- | --- | --- | --- | --- |
| **Age (median cal. BP)** |  | 617 | 1613 | 2892 | 3843 | 4658 | 5006 | 5177 | 5480 | 5923 | 6935 | 7458 | 8296 | 9022 | 9407 | 9808 |
| **Pollen concentration** |  | High | High | High | High | High | High | High | High | High | High | High | High | High | Low-medium | Very low |
| **Preservation** |  | Medium | Low-medium | Medium-low | Low | Low | Low | Low | Medium | Medium-low | Low-medium | Low-medium | Low-medium | Medium | Medium | Low |
| Trees | **Species** |  | | | | | | | | | | | | | | |
|  | Alnus | 10.41 | 13.88 | 12.83 | 20.66 | 16.52 | 23.80 | 16.73 | 19.38 | 25.36 | 28.76 | 31.76 | 27.75 | 1.29 | 0.00 | 1.01 |
|  | Betula | 24.66 | 29.99 | 25.95 | 33.53 | 31.23 | 35.63 | 38.23 | 42.98 | 40.29 | 37.69 | 30.88 | 21.98 | 29.90 | 42.79 | 27.27 |
|  | Carpinus | 0.31 | 0.00 | 0.00 | 0.00 | 0.00 | 0.00 | 0.00 | 0.00 | 0.00 | 0.00 | 0.00 | 0.00 | 0.00 | 0.00 | 0.00 |
|  | Corylus | 1.53 | 3.99 | 2.04 | 2.60 | 6.16 | 4.79 | 2.04 | 3.51 | 7.13 | 5.45 | 4.12 | 5.22 | 3.05 | 3.72 | 3.03 |
|  | Fraxinus | 0.31 | 0.44 | 0.29 | 0.43 | 0.00 | 0.30 | 0.54 | 0.42 | 0.00 | 0.00 | 0.00 | 0.00 | 0.00 | 0.00 | 0.00 |
|  | Picea | 10.11 | 0.15 | 0.00 | 0.00 | 0.00 | 0.00 | 0.00 | 0.00 | 0.00 | 0.00 | 0.00 | 0.00 | 0.00 | 0.00 | 0.00 |
|  | Pinus | 33.23 | 34.42 | 47.67 | 26.73 | 31.98 | 18.41 | 27.35 | 20.51 | 12.81 | 13.39 | 17.50 | 30.77 | 57.07 | 32.09 | 15.15 |
|  | Quercus | 0.92 | 1.48 | 0.73 | 1.30 | 1.65 | 0.30 | 0.82 | 1.54 | 0.40 | 0.17 | 0.29 | 0.55 | 0.64 | 0.93 | 0.00 |
|  | Tilia | 0.00 | 0.30 | 0.00 | 0.14 | 0.45 | 0.30 | 0.27 | 0.56 | 0.13 | 0.33 | 0.15 | 0.00 | 0.00 | 0.00 | 0.00 |
|  | Ulmus | 0.61 | 0.59 | 1.31 | 0.72 | 0.30 | 0.60 | 1.09 | 1.40 | 1.98 | 2.81 | 2.21 | 0.96 | 0.32 | 0.16 | 0.00 |
|  | Populus | 0.15 | 0.00 | 0.15 | 0.58 | 0.15 | 0.15 | 0.00 | 0.14 | 0.00 | 0.00 | 0.15 | 0.00 | 0.00 | 0.16 | 0.00 |
| Shrubs | Hippophae | 0.00 | 0.00 | 0.00 | 0.00 | 0.00 | 0.15 | 0.00 | 0.00 | 0.00 | 0.00 | 0.00 | 0.00 | 0.00 | 1.71 | 2.02 |
|  | Juniperus | 0.46 | 0.89 | 0.15 | 0.58 | 0.15 | 0.30 | 0.41 | 0.42 | 0.00 | 0.33 | 0.15 | 0.14 | 0.16 | 0.62 | 1.01 |
|  | Myrica | 0.00 | 0.00 | 0.00 | 0.00 | 0.00 | 0.00 | 0.14 | 0.14 | 0.13 | 0.00 | 0.15 | 0.14 | 0.00 | 0.00 | 0.00 |
|  | Salix | 1.53 | 2.07 | 0.29 | 1.01 | 0.45 | 0.90 | 0.00 | 0.56 | 0.92 | 0.66 | 1.18 | 2.06 | 0.64 | 5.74 | 2.02 |
|  | Frangula alnus | 0.00 | 0.00 | 0.00 | 0.00 | 0.00 | 0.00 | 0.00 | 0.00 | 0.00 | 0.17 | 0.00 | 0.00 | 0.00 | 0.00 | 0.00 |
|  | Sorbus | 0.46 | 0.15 | 0.15 | 1.16 | 0.30 | 0.45 | 0.14 | 0.56 | 0.66 | 0.33 | 0.74 | 0.55 | 0.16 | 0.00 | 0.00 |
|  | Calluna | 0.15 | 0.15 | 0.15 | 0.29 | 0.00 | 0.30 | 0.00 | 0.14 | 0.26 | 0.00 | 0.00 | 0.00 | 0.00 | 0.00 | 0.00 |
|  | Empetrum | 0.00 | 0.00 | 0.00 | 0.00 | 0.00 | 0.00 | 0.00 | 0.00 | 0.00 | 0.00 | 0.00 | 0.00 | 0.16 | 0.16 | 0.00 |
|  | Ericaceae undiff | 0.15 | 0.15 | 0.00 | 0.00 | 0.15 | 0.00 | 0.14 | 0.14 | 0.13 | 0.00 | 0.00 | 0.00 | 0.16 | 0.31 | 0.00 |
|  | Cf. Prunus padus | 0.00 | 0.00 | 0.00 | 0.00 | 0.00 | 0.00 | 0.00 | 0.00 | 0.13 | 0.17 | 0.00 | 0.14 | 0.00 | 0.00 | 0.00 |
| Cultivated | Avena | 0.00 | 0.00 | 0.00 | 0.00 | 0.00 | 0.00 | 0.00 | 0.00 | 0.00 | 0.00 | 0.00 | 0.14 | 0.00 | 0.00 | 0.00 |
|  | Hordeum | 0.15 | 0.00 | 0.00 | 0.00 | 0.00 | 0.00 | 0.00 | 0.00 | 0.00 | 0.00 | 0.00 | 0.00 | 0.00 | 0.00 | 0.00 |
| Herbs | Cannabis type | 0.31 | 0.30 | 0.15 | 0.14 | 0.00 | 0.15 | 0.14 | 0.42 | 0.00 | 0.17 | 0.00 | 0.00 | 0.00 | 0.16 | 0.00 |
|  | Humulus | 0.00 | 0.30 | 0.15 | 0.43 | 0.00 | 0.15 | 0.00 | 0.00 | 0.00 | 0.00 | 0.00 | 0.00 | 0.00 | 0.00 | 0.00 |
|  | Poaceae | 5.05 | 2.07 | 0.87 | 0.43 | 0.45 | 0.45 | 0.41 | 0.84 | 1.45 | 0.66 | 2.79 | 0.55 | 2.25 | 2.48 | 6.06 |
|  | Cyperaceae | 1.84 | 0.74 | 0.44 | 1.01 | 0.90 | 2.40 | 1.09 | 2.25 | 2.38 | 1.65 | 1.32 | 2.88 | 0.64 | 1.86 | 3.03 |
|  | Apiaceae | 0.00 | 0.15 | 0.00 | 0.00 | 0.30 | 0.00 | 0.00 | 0.00 | 0.00 | 0.00 | 0.00 | 0.00 | 0.16 | 0.00 | 1.01 |
|  | Artemisia | 0.15 | 0.15 | 0.15 | 0.00 | 0.15 | 0.00 | 0.14 | 0.00 | 0.00 | 0.00 | 0.00 | 0.14 | 0.00 | 0.00 | 0.00 |
|  | Crepis-t | 0.00 | 0.00 | 0.00 | 0.14 | 0.15 | 0.00 | 0.14 | 0.00 | 0.00 | 0.00 | 0.00 | 0.00 | 0.00 | 0.00 | 0.00 |
|  | Senecio-type | 0.31 | 0.00 | 0.00 | 0.14 | 0.00 | 0.00 | 0.14 | 0.00 | 0.00 | 0.00 | 0.00 | 0.14 | 0.00 | 0.00 | 0.00 |
|  | Valeriana sambucifolia | 0.00 | 0.00 | 0.15 | 0.14 | 0.00 | 0.15 | 0.00 | 0.00 | 0.00 | 0.00 | 0.00 | 0.00 | 0.00 | 0.00 | 0.00 |
|  | Epilobium | 0.00 | 0.00 | 0.15 | 0.00 | 0.00 | 0.00 | 0.00 | 0.14 | 0.00 | 0.00 | 0.00 | 0.00 | 0.00 | 0.16 | 0.00 |
|  | Cerastium-t | 0.00 | 0.00 | 0.29 | 0.00 | 0.00 | 0.15 | 0.00 | 0.00 | 0.00 | 0.00 | 0.00 | 0.00 | 0.00 | 0.00 | 0.00 |
|  | Melampyrum | 0.00 | 0.00 | 0.00 | 0.00 | 0.00 | 0.15 | 0.00 | 0.00 | 0.00 | 0.00 | 0.00 | 0.14 | 0.16 | 0.00 | 0.00 |
|  | Chenopodiaceae | 0.15 | 0.00 | 0.00 | 0.00 | 0.00 | 0.00 | 0.00 | 0.00 | 0.00 | 0.17 | 0.00 | 0.00 | 0.00 | 0.00 | 0.00 |
|  | Chenopodium | 0.00 | 0.00 | 0.00 | 0.14 | 0.00 | 0.00 | 0.00 | 0.00 | 0.00 | 0.00 | 0.00 | 0.00 | 0.00 | 0.00 | 0.00 |
|  | Plantago major-med.-t | 0.00 | 0.15 | 0.00 | 0.00 | 0.00 | 0.00 | 0.00 | 0.00 | 0.00 | 0.00 | 0.00 | 0.00 | 0.00 | 0.00 | 0.00 |
|  | Filipendula | 1.53 | 1.62 | 0.29 | 1.45 | 0.15 | 1.95 | 0.82 | 0.42 | 1.19 | 0.17 | 0.29 | 0.14 | 0.32 | 0.16 | 0.00 |
|  | Galium | 0.00 | 0.00 | 0.00 | 0.00 | 0.00 | 0.00 | 0.14 | 0.00 | 0.00 | 0.00 | 0.00 | 0.00 | 0.00 | 0.00 | 0.00 |
|  | Polygonum aviculare | 0.15 | 0.15 | 0.00 | 0.00 | 0.00 | 0.00 | 0.00 | 0.00 | 0.00 | 0.00 | 0.00 | 0.00 | 0.00 | 0.00 | 0.00 |
|  | Potentilla-t | 0.00 | 0.00 | 0.00 | 0.00 | 0.00 | 0.15 | 0.00 | 0.00 | 0.00 | 0.00 | 0.00 | 0.00 | 0.00 | 0.00 | 0.00 |
|  | Ranunculaceae | 0.00 | 0.00 | 0.00 | 0.00 | 0.15 | 0.00 | 0.00 | 0.14 | 0.00 | 0.00 | 0.15 | 0.00 | 0.00 | 0.00 | 0.00 |
|  | Ranunculus acris type | 0.00 | 0.00 | 0.15 | 0.00 | 0.00 | 0.00 | 0.00 | 0.00 | 0.00 | 0.00 | 0.15 | 0.14 | 0.00 | 0.00 | 0.00 |
|  | Rosaceae undiff | 0.00 | 0.00 | 0.00 | 0.43 | 0.00 | 0.15 | 0.00 | 0.00 | 0.00 | 0.00 | 0.00 | 0.00 | 0.00 | 0.00 | 0.00 |
|  | Rumex | 0.46 | 0.15 | 0.44 | 0.29 | 0.00 | 0.15 | 0.00 | 0.00 | 0.00 | 0.00 | 0.00 | 0.27 | 0.00 | 0.16 | 1.01 |
|  | Urtica | 0.46 | 0.00 | 0.00 | 0.00 | 0.00 | 0.00 | 0.00 | 0.00 | 0.00 | 0.00 | 0.00 | 0.00 | 0.00 | 0.00 | 1.01 |
| Vascular cryptogams/mosses | Selaginella | 0.15 | 0.00 | 0.00 | 0.00 | 0.00 | 0.00 | 0.00 | 0.00 | 0.00 | 0.00 | 0.00 | 0.00 | 0.00 | 0.00 | 0.00 |
|  | Equisetum | 0.00 | 0.15 | 0.15 | 1.01 | 0.00 | 0.00 | 0.27 | 0.14 | 0.13 | 0.00 | 0.00 | 0.00 | 0.00 | 0.16 | 1.01 |
|  | Polypodiaceae undiff | 12.10 | 9.60 | 12.24 | 11.56 | 14.41 | 30.24 | 18.78 | 17.28 | 15.98 | 12.07 | 12.06 | 4.26 | 1.93 | 15.35 | 4.04 |
|  | Sphagnum | 0.61 | 0.00 | 0.00 | 0.14 | 0.00 | 0.15 | 0.14 | 0.14 | 0.00 | 0.00 | 0.00 | 0.00 | 0.16 | 0.93 | 1.01 |
|  | Huperzia selago | 0.00 | 0.00 | 0.00 | 0.00 | 0.00 | 0.00 | 0.00 | 0.00 | 0.00 | 0.17 | 0.00 | 0.00 | 0.00 | 0.00 | 0.00 |
|  | Lycopodium sp | 0.00 | 0.44 | 0.29 | 0.72 | 0.00 | 0.00 | 0.82 | 0.28 | 0.26 | 0.17 | 0.15 | 0.14 | 0.00 | 0.00 | 1.01 |
| Aquatic plants | Myriophyllum alterniflorum | 0.00 | 0.30 | 0.15 | 0.00 | 0.00 | 0.00 | 0.00 | 0.00 | 0.00 | 0.00 | 0.29 | 0.14 | 0.00 | 0.00 | 0.00 |
|  | Sparganium-t | 0.00 | 0.00 | 0.00 | 0.14 | 0.00 | 0.30 | 0.00 | 0.14 | 0.00 | 0.00 | 0.00 | 0.00 | 0.00 | 0.00 | 0.00 |
|  | Menyanthes trifoliata | 0.00 | 0.00 | 0.00 | 0.00 | 0.00 | 0.15 | 0.00 | 0.14 | 0.00 | 0.17 | 0.00 | 0.14 | 0.32 | 0.31 | 0.00 |
|  | Lemna cf minor | 0.15 | 0.00 | 0.29 | 0.00 | 0.00 | 0.00 | 0.00 | 0.00 | 0.00 | 0.00 | 0.00 | 0.00 | 0.00 | 0.00 | 0.00 |
|  | Isoetes echinospore | 2.14 | 5.17 | 4.52 | 2.75 | 2.25 | 2.69 | 1.50 | 3.37 | 2.77 | 0.83 | 0.59 | 0.82 | 0.64 | 0.00 | 0.00 |
| Other | Pediastrum | 0.00 | 0.15 | 0.15 | 0.00 | 0.00 | 0.00 | 0.00 | 0.00 | 0.00 | 0.00 | 0.00 | 0.27 | 0.00 | 0.16 | 0.00 |
|  | Botryococcus | 0.31 | 0.30 | 0.00 | 0.72 | 0.45 | 0.30 | 0.00 | 0.42 | 0.66 | 0.33 | 0.00 | 0.00 | 0.00 | 0.00 | 0.00 |
|  | Ephedra distachya-t | 0.00 | 0.00 | 0.00 | 0.00 | 0.00 | 0.00 | 0.00 | 0.00 | 0.00 | 0.00 | 0.00 | 0.00 | 0.00 | 0.16 | 0.00 |
|  | Charcoal > 20cm | 2.45 | 1.62 | 0.29 | 1.16 | 1.50 | 3.59 | 4.22 | 1.97 | 2.11 | 2.81 | 2.79 | 1.65 | 2.73 | 14.26 | 202.02 |
|  | Indet. pollen | 4.44 | 5.61 | 5.10 | 5.49 | 8.26 | 7.63 | 9.12 | 3.37 | 4.62 | 6.94 | 6.03 | 5.22 | 2.89 | 6.67 | 36.36 |
|  | **Pollensum** | **653** | **677** | **686** | **692** | **666** | **668** | **735** | **712** | **757** | **605** | **680** | **728** | **622** | **645** | **99** |

*Table S7. Pollen data from South Mesna lake. The division between Cannabis-type and Humulus is explained in comments below*

*Low magnification analysis*

Eight types of pollen/spores were found and identified in the analysis (Table S8). They relate to a larger pollen sum (between 7000 and 21,000 pollen), and their percentage value has been calculated with respect to this. The percentage value is of course of little importance, but the finding of pollen itself is most important, for example rye pollen was found at 10 cm, which was not found in the standard analysis, and thus it is likely that we can conclude that rye cultivation occurred in the vicinity around the 16th century. The earliest finding of spruce pollen in the standard analysis was at 30 cm, but the low-magnification analysis found a pollen already at the 90 cm level (ca. 4000 cal. BP). It seems likely that in this case it is long-flight pollen, perhaps spread from spruce trees northeast of the mountain range. Pollen of beech (*Fagus sylvatica*) was also found at 70 cm (3400 cal. BP), which is a very early find. Beech pollen was also found at 10 cm (16th century). Here it is possible that it is a question of local/extra-local presence of beech at the mentioned time periods. For example, a stand of beech may have existed in the valley at Mjøsa, or even at the Mesna lakes.

| ***Depth*** | | **10** | **30** | **50** | **70** | **90** | **110** | **120** | **140** | **160** | **180** | **200** | **230** | **250** | **270** | **290** |
| --- | --- | --- | --- | --- | --- | --- | --- | --- | --- | --- | --- | --- | --- | --- | --- | --- |
|  | Picea | 7.831 | 0.047 | 0.000 | 0.012 | 0.008 | 0.000 | 0.000 | 0.000 | 0.000 | 0.000 | 0.000 | 0.000 | 0.000 | 0.000 | 0.000 |
|  | Fagus | 0.012 | 0.000 | 0.000 | 0.024 | 0.000 | 0.000 | 0.000 | 0.000 | 0.000 | 0.000 | 0.000 | 0.000 | 0.000 | 0.000 | 0.000 |
|  | Tilia | 0.024 | 0.000 | 0.000 | 0.000 | 0.055 | 0.000 | 0.000 | 0.000 | 0.000 | 0.000 | 0.016 | 0.000 | 0.000 | 0.000 | 0.000 |
|  | Carpinus | 0.048 | 0.000 | 0.000 | 0.000 | 0.000 | 0.000 | 0.000 | 0.000 | 0.000 | 0.000 | 0.000 | 0.000 | 0.000 | 0.000 | 0.000 |
|  | Secale | 0.024 | 0.000 | 0.000 | 0.000 | 0.000 | 0.000 | 0.000 | 0.000 | 0.000 | 0.000 | 0.000 | 0.000 | 0.000 | 0.000 | 0.000 |
|  | Cerealia | 0.024 | 0.000 | 0.000 | 0.000 | 0.000 | 0.000 | 0.000 | 0.000 | 0.000 | 0.000 | 0.000 | 0.000 | 0.000 | 0.000 | 0.000 |
|  | Valeriana sambucifolia | 0.000 | 0.000 | 0.008 | 0.000 | 0.024 | 0.008 | 0.000 | 0.000 | 0.000 | 0.000 | 0.000 | 0.000 | 0.000 | 0.000 | 0.000 |
|  | Dryopteris f-mas | 0.000 | 0.000 | 0.000 | 0.000 | 0.000 | 0.016 | 0.000 | 0.000 | 0.000 | 0.039 | 0.008 | 0.000 | 0.000 | 0.000 | 0.000 |
|  | Picea | 7.831 | 0.047 | 0.000 | 0.012 | 0.008 | 0.000 | 0.000 | 0.000 | 0.000 | 0.000 | 0.000 | 0.000 | 0.000 | 0.000 | 0.000 |
|  | Fagus | 0.012 | 0.000 | 0.000 | 0.024 | 0.000 | 0.000 | 0.000 | 0.000 | 0.000 | 0.000 | 0.000 | 0.000 | 0.000 | 0.000 | 0.000 |
|  | Tilia | 0.024 | 0.000 | 0.000 | 0.000 | 0.055 | 0.000 | 0.000 | 0.000 | 0.000 | 0.000 | 0.016 | 0.000 | 0.000 | 0.000 | 0.000 |
|  | 100xMAG pollen sum | **8300** | **12900** | **13000** | **8500** | **12700** | **12700** | **9300** | **13500** | **9600** | **7700** | **12900** | **13800** | **21000** | **4100** | **400** |

*Table S8. Results from a low magnification analysis (100 x analysis) presented in percentages. It has been conducted separately to detect low-frequency pollen taxa, mainly cultivated grain. They therefore relate to a larger pollen sum, which here is the total interpolated pollen sum of one or two microscope slides.*

*South Mesna lake as an archive for pollen and its catchment area*

The lake is a regional pollen sink, i.e. the pollen composition in the sediments encapsulate a large geographic area, many kilometers from the lake. This applies especially to trees, such as pine, birch and hazel but also other wind-pollinated plants, and to a lesser extent shrubs, herbs and insect-pollinated plants. The fact that the pollen collection area is so large (cf. (32)), makes parts of the vegetation interpretation complicated. For example, tree pollen probably comes from the "region", while pollen from herbs and insect-pollinated plants mainly comes from the immediate area, i.e. the slopes of the valley and the drainage area itself. There are few modern examples of published pollen diagrams from such large lake locations to compare with, most pollen analytical studies from Norway and Sweden are carried out on small lakes or peatlands (cf. (33-35)).

The pollen diagram from South Mesna has been divided into five periods (Figure S6), or pollen zones, based on interpretation of all available pollen data (Table S7). The zoning is thus based on changes in the entire pollen composition, all data has been considered.

**Period A (approx. 9800 – 9300 cal. BP).** The sample is from the time of the deglaciation or shortly after. Possibly, melting permafrost may still have existed in the valley or in the drainage area. Light-demanding plants dominate locally: birch, hazel, alder, willow/willow and herbs such as grasses, sedges and sorrels. Heather plants such as crowberry/crickling (*Empetrum* sp.) appear to grow within the drainage area. It is also likely that oak immigrates to the immediate area during this time, as well as elm and aspen. Spores from ferns (family Polypodiaceae) also occur, they grow locally in the immediate area. The pollen content of sea buckthorn/buckthorn (*Hippophae* sp.) reaches up to 2% and the shrub probably grows locally in the valley. Around this time, or slightly earlier, the marine shoreline is probably at Mjøsa/Lillehammer, and this bush follows the displacement of the beach. Also, a pollen grain of *Ephedra distachya* was found in the second lowest level at 270 cm (ca. 9400 cal BP), it is probably redeposited from late glacial sediments (cf. (35)). *Ephedra* is a genus of leafless shrubs that, among other things, occur in the Mediterranean area and on the Ukrainian steppes today. It is still unclear whether it grew "locally" in Scandinavia during the Late Glacial, but it seems likely. The content of charcoal is higher during this period than later, they have accumulated in the drainage area for a long time and are now concentrated in the basin.

**Period B (approx. 9300 – 8500 cal. BP).** Shortly after the beginning of the period, pine reaches its maximum percentage value, which may reflect the regional dominance of pine during the period. Percentage values for almost all other taxa decrease as a result, which does not necessarily mean that they decrease in the local vegetation. Rowan (*Sorbus aucuparia*) is increasing, however, probably in the immediate area. Pollen of *Filipendula* sp. (probably meadowsweet) (appears at this time, probably growing along the shores of the lake. It occurs here before alder has expanded locally, which has been observed in several previous surveys. At the end of the period, alder tree pollen significantly increases, reflecting a local expansion of alder (*Alnus glutinosa)* along the shores of the lake. Periods A and B are similar to pollen zone H1 - H4 from Hemma (36), and are partly deposited during the same time, but in completely different environments.

**Period C (8500 – 5800 cal. BP).** At the beginning of the period, alder becomes dominant in some places along the shores of the lakes. Hazel gradually increases towards its highest values at the end of the period, which possibly also reflects the local presence. Alm increases during the beginning of the period and reaches its highest value around 7000 cal. BP. Linden also took place in the local landscape in the middle of the period. At the end of the period, there is also occasional pollen from heather (*Calluna* sp.), the light-demanding shrub may then have started to expand in local altitude areas. Wetlands with semi-grass and possibly willow and elk grass probably increase in the drainage area throughout the period. Pollen from mugworts, acids and mollusks appear around 8300 cal BP, and although they may have grown in the coastal zones, it is very uncertain if they reflect human activity, such as Mesolithic settlements along the beaches. From the same time there is an Avena-type pollen, it is probably flying oats (*Avena fatua*), although the pollen grain is almost identical to cultivated oats. Wild oat is common as a weed in cultivated fields but occurs here perhaps in the beach zone. Bird cherry (*Prunus padus*), an insect-pollinated tree with edible fruit, also grows in the valley during this period.

**Period D (5800 – 1300 cal. BP).** The period begins with hazel decreasing significantly. Oak and linden increase clearly at the beginning of the period, and ash pollen appears for the first time. Pollen from ash (*Fraxinus excelsior*) can be problematic to determine due to poor pollen preservation, but several better-preserved grains were found, starting at 70 cm (ca. 5500 cal. BP). Ash is wind-pollinated and can spread relatively far but has probably grown in the valley at South Mesna already in earlier periods. It seems unlikely that the ash pollen has spread here from, for example, the Mjøsa area in the west. There is no clear increase in light-demanding herbs during the beginning of the period, so it seems unlikely that any pastures are established at this time. Alder also clearly decreases from the beginning of the period, reaching a minimum value around 5000 cal. BP. It is possible that this has to do with human activity on the beaches, maybe alder is being cleared from certain places, for example there are weeds such as mugwort occurring at the same time. Juniper (*Juniperus* sp.) increases at the same time, but it is possible we see changes in the vegetation on different soil types or different parts of the valley at the same time. In contrast, between about 5200 and 4600 cal. BP there is a change in the charcoal curve, accompanied by clear increases in the Polypodiaceae, meadowsweet and sedge curves. A contemporary decline in the LOI curve likely indicates increased soil erosion in the drainage area. At this time, there is also no clear change in herbs and grasses, but all deciduous trees decrease, which could mean that local forest grazing occurs. Heather also increases slightly at this time, possibly associated with the use of clearing fires, which can explain the increase in charcoal. Around 3400 cal. BC and beyond, sedges, grasses and heather seem to increase slightly while species such as linden decrease in the immediate area, this is probably a consequence of herbaceous pastures becoming more permanent features of the valley. Around 1600 cal. BP juniper reaches its maximum value, which probably is associated with the spread of pastures.

**Period E (600 – 500 cal. BP).** The period consists of only one sample, but contains clear traces of, among other things, cultivation and pastures. Acids and grass reach their highest values at this time, which probably indicates that the most extensive pastures existed during this period, although the percentage values are still very modest. It is unclear why certain classic pasture-indicative herbs are completely missing from the pollen spectrum, for example black sedge (*Plantago lanceolata*). Pollen from barley (*Hordeum* sp.) is found in the top sample and indicates local cultivation and threshing sites in the valley near South Mesna. Pollen from rye was also found in the low-magnification analysis (Figure S8), and it is possible that it is autumn-sown rye, which became significantly more common in Scandinavia during the 16th century. Spruce expands strongly at this time and reaches around 10%, perhaps colonizing abandoned fields and pastures in the wake of the black death (e.g. (37)). However, it is difficult to determine whether the spruce's expansion is truly local or regional. Other tree species decrease slightly, although pine, birch and oak maintain close to stable values. Although hazel has decreased somewhat, it remains in the immediate area. Hornbeam (Carpinus betulus) pollen occurs at this time, as does beech (see the low-magnification analysis above), indicating local presence of these tree species. Hornbeam may have been a technologically important tree species, as the wood is the heaviest and hardest found in Scandinavia.

*Comments regarding Cannabis-type (Cannabinaceae) and Humulus*

The division is based on measurements of the diameter of the pollen grains according to Beug (31). Since no measurements resulted in a determination of hemp (*Cannabis sativa*), it is likely that all Cannabis-type pollen grains found in the study actually come from hops (*Humulus lupulus*). It seems most likely that these are wild hops, as the percentages are very low throughout the sequence. However, hops have probably still been cultivated in the area, for example during the late Middle Ages and later.

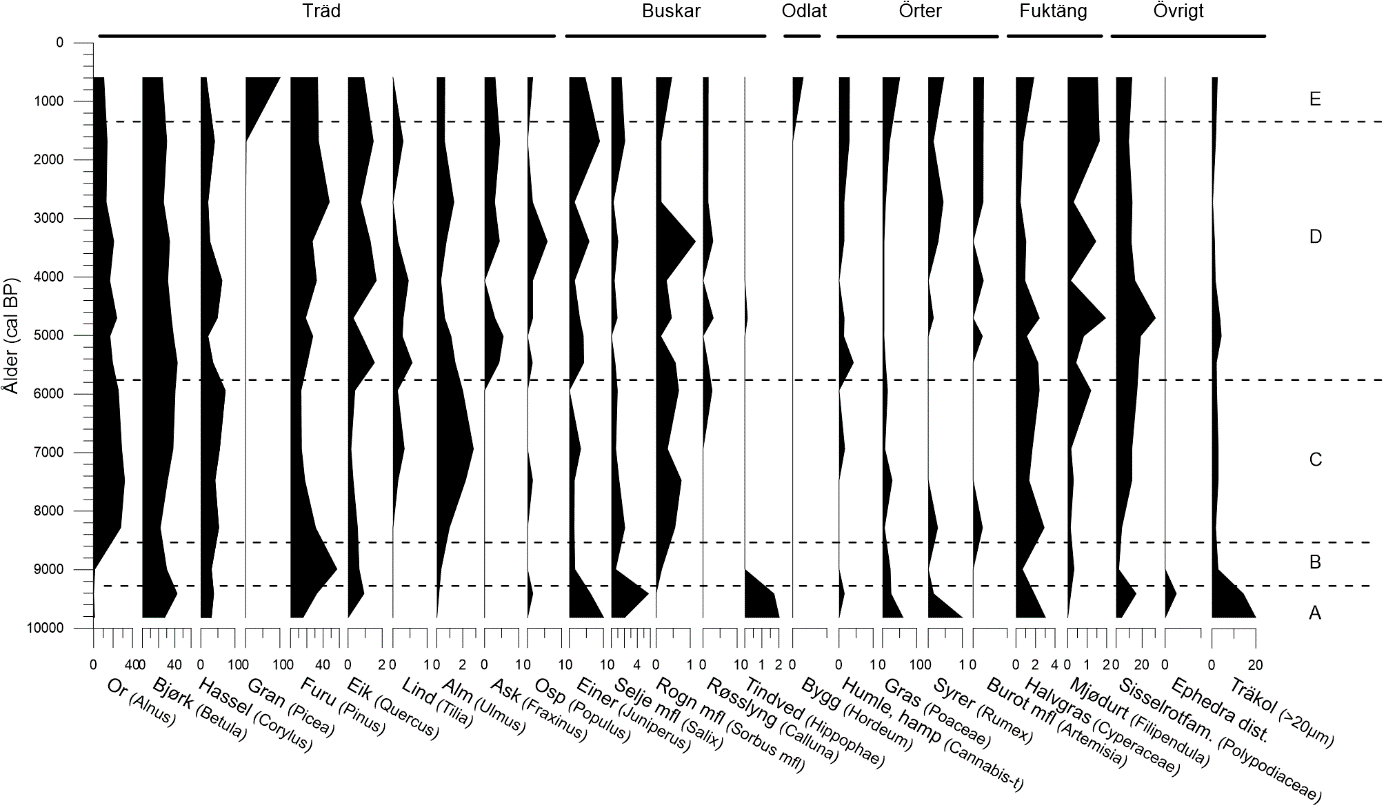

*Figure S7. Pollen diagram from South Mesna lake. Percentage data from the most important pollen types. The diagram is divided into five periods/pollen zones.*

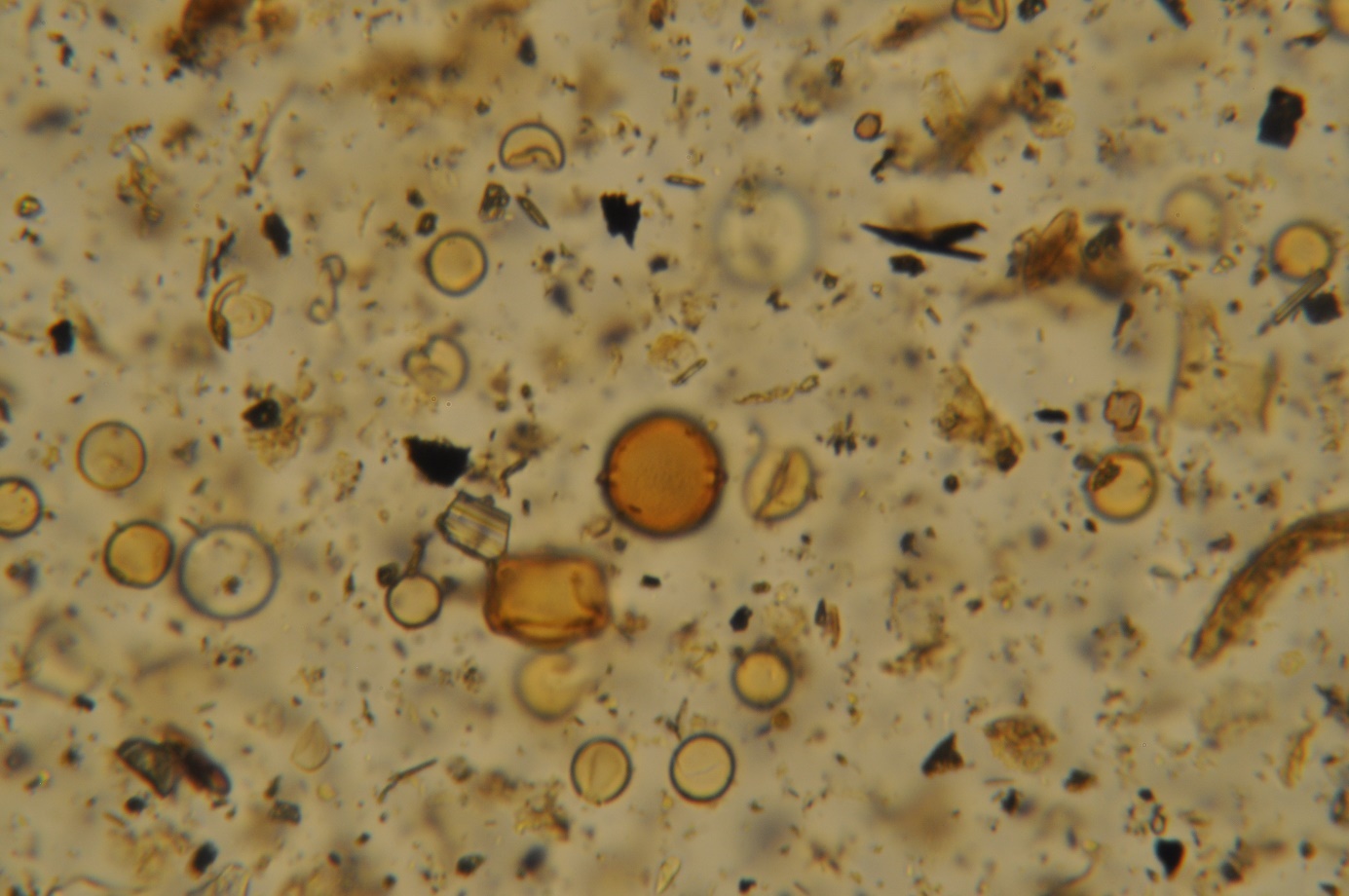

Figure S8. Pollen from sea buckthorn (Hippophaë rhamnoides).

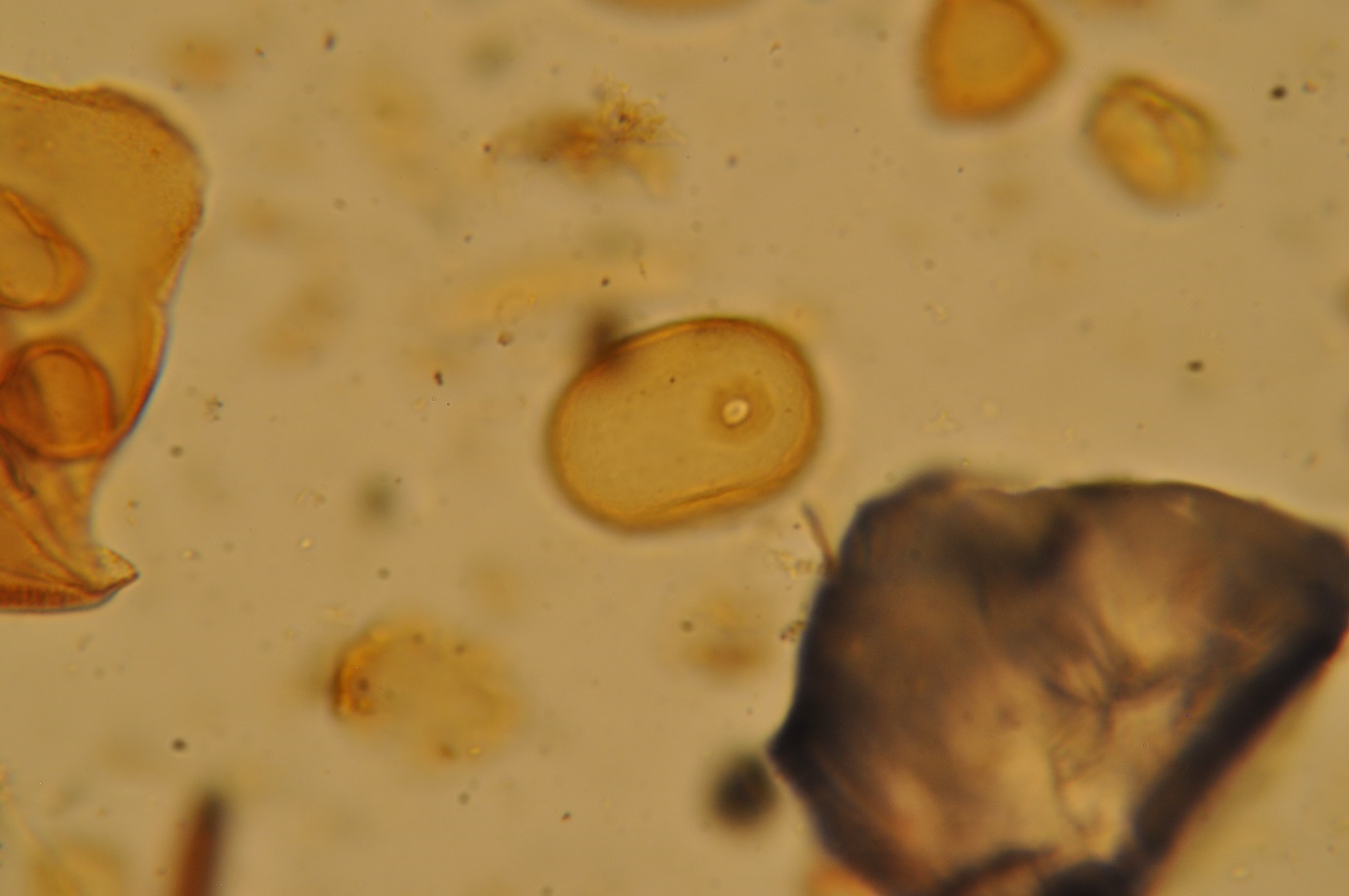

Figure S9. Pollen from rye (Secale cereale).
